# Exploring endothelial cell environments across organs in spatially resolved omics data

**DOI:** 10.1101/2025.09.23.678129

**Authors:** Yashvardhan Jain, Jodie Jepson, Roy Chen, Elizabeth Maier, Bruce W. Herr, Aleix Puig-Barbe, Ellen M. Quardokus, Danial Qaurooni, Clarence Yapp, Samuel L. Ewing, Archibald Enninful, Negin Farzad, Andreas Bueckle, Quinn T. Easter, Bruno Matuck, Chenchen Zhu, Emma Marie Monte, Jeffrey M. Purkerson, Matthew Jehrio, Ravi S. Misra, Rong Fan, Fiona Ginty, Arivarasan Karunamurthy, Jean Fan, Martha Campbell-Thompson, Gloria S. Pryhuber, Kevin M. Byrd, John W. Hickey, Katy Börner

**Affiliations:** Department of Intelligent Systems Engineering, Luddy School of Informatics, Computing, and Engineering, Indiana University, Bloomington, IN 47408, USA; Department of Biomedical Engineering, Duke University, Durham, NC, USA; European Bioinformatics Institute (EMBL-EBI), Wellcome Genome Campus, Hinxton, Cambridge CB10 1SD, UK; Laboratory of Systems Pharmacology, Harvard Medical School, Boston, MA, USA; Department of Pathology, Immunology and Laboratory Medicine, College of Medicine, University of Florida, Gainesville FL 32610 USA; Department of Biomedical Engineering, Yale University, New Haven, CT, USA; Department of Oral and Craniofacial Molecular Biology, Virginia Commonwealth University, Richmond, VA, USA; Department of Genetics, Stanford School of Medicine, Stanford, CA 94305; University of Rochester Medical Center, Rochester, NY, USA; GE HealthCare Technology and Innovation Center, Niskayuna, NY 12309; Department of Dermatology & Pathology, University of Pittsburgh Medical Center, Pittsburgh, PA 15213, USA; Department of Biomedical Engineering, Johns Hopkins University, Baltimore, MD, USA

## Abstract

Endothelial cells are ubiquitously present in the human body and line the luminal surface of blood and lymphatic vessels. The oxygen-dependence of cells impacts their proximity to blood vessels, and consequently, to endothelial cells depending on their functional properties and priorities. This paper presents cell-to-nearest-endothelial-cell distance distributions for various cell types using 399 spatially resolved omics datasets from 14 studies comprising 12 tissue types with a total of 47,349,496 cells. Additionally, we developed an open-source web-based interactive tool, Cell Distance Explorer, that allows researchers to interactively visualize cell graphs and linkages in 2D and 3D datasets. Finally, we present a hierarchical neighborhood analysis focused on the endothelial cell neighborhoods in small and large intestine datasets. This paper provides an open-access resource (datasets, tools, and analyses) to characterize and compare cell distances and cell neighborhoods in spatially resolved omics data.

## Introduction

*Endothelial Cells (ECs)* line the luminal surface of blood and lymphatic vessels, serving as a barrier and regulating exchanges between blood and lymphatic circulation and the surrounding tissue^1^. The vasculature forms an uninterrupted path across scales in the human body–connecting cells, functional tissue units^2^, and organs, and enabling the flow of oxygen, nutrients, and drugs throughout the human body.

Various cell atlassing efforts^3–10^ have been working to further our understanding of cellular organization and relationships, especially with the advances in spatially resolved omics technologies–spatial transcriptomics and spatial proteomics–which enable high resolution single-cell molecular profiling to identify distinct cell types while preserving the spatial organization of cells in the tissue^11–13^. Other efforts have focused on building an open, computer-readable, and comprehensive database of the adult human blood vasculature to create a better common coordinate framework using the vasculature^2,14,15^, and understanding the angiodiversity and organotypic molecular signatures in human vascular cells^5^.

The oxygen-dependence of all cells impacts their proximity to ubiquitously present *ECs*, thereby impacting cellular organization in different tissue regions and organs based on the functional and structural properties and priorities of different cell populations. For example, immune cells might generally have lower distances to endothelial cells due to their need for rapid surveillance and circulation. Given spatially resolved omics data, it becomes possible to analyze cellular networks and evaluate the spatial relationships between different cell populations relative to blood vasculature and further our understanding of the role of human blood vasculature in spatial organization of cells across organs, tissue regions, and conditions. Specifically, we can focus on absolute distances between cells to analyze and visualize spatial networks of cells in various tissues and compare the emergent patterns in the distance distributions for different cell types and their nearest endothelial cells.

In this paper, we conduct a meta-analysis of publicly available spatially resolved omics data by curating and cataloging the endothelial cell populations and cell-to-nearest-endothelial-cell distance distributions for various cell types across tissues. We collect, analyze, and visualize cell networks using a variety of datasets from different tissues, both normal and diseased, and 2D and 3D. Previously developed tools–Registration User Interface^16^ and Exploration User Interface^16^–are used to spatially register and catalog the tissue specimen origin in a common 3D framework of the Human Reference Atlas^6^ (https://humanatlas.io). We unify the cell type labels from data providers into a 3-level typology, including mapping each cell type to Cell Ontology. Additionally, we demonstrate the value of *ECs* within cellular neighborhoods, revealing cell-to-nearest-endothelial-cell interactions within hierarchical cellular neighborhoods^17^ in the intestine.

We also present an open-source web-based tool for researchers, Cell Distance Explorer (see **Fig. 1a**), that allows uploading such datasets to compute and visualize cell-to-nearest-anchor-cell linkages and distance distributions in an interactive environment, where an anchor cell is defined as a cell type of interest the nearest distance (Euclidean) to which is calculated from all other cell types in its proximity. This free-of-cost tool can benefit researchers who are interested in analyzing and comparing the changes in cell-cell distance distributions between normal and disease tissue, across diseases, and between different tissue regions to evaluate how different conditions impact cell organization in tissues. All datasets, tools, and analyses detailed in this paper are provided as an open-access resource, see **Data** and **Code Availability**.

**Figure 1.**
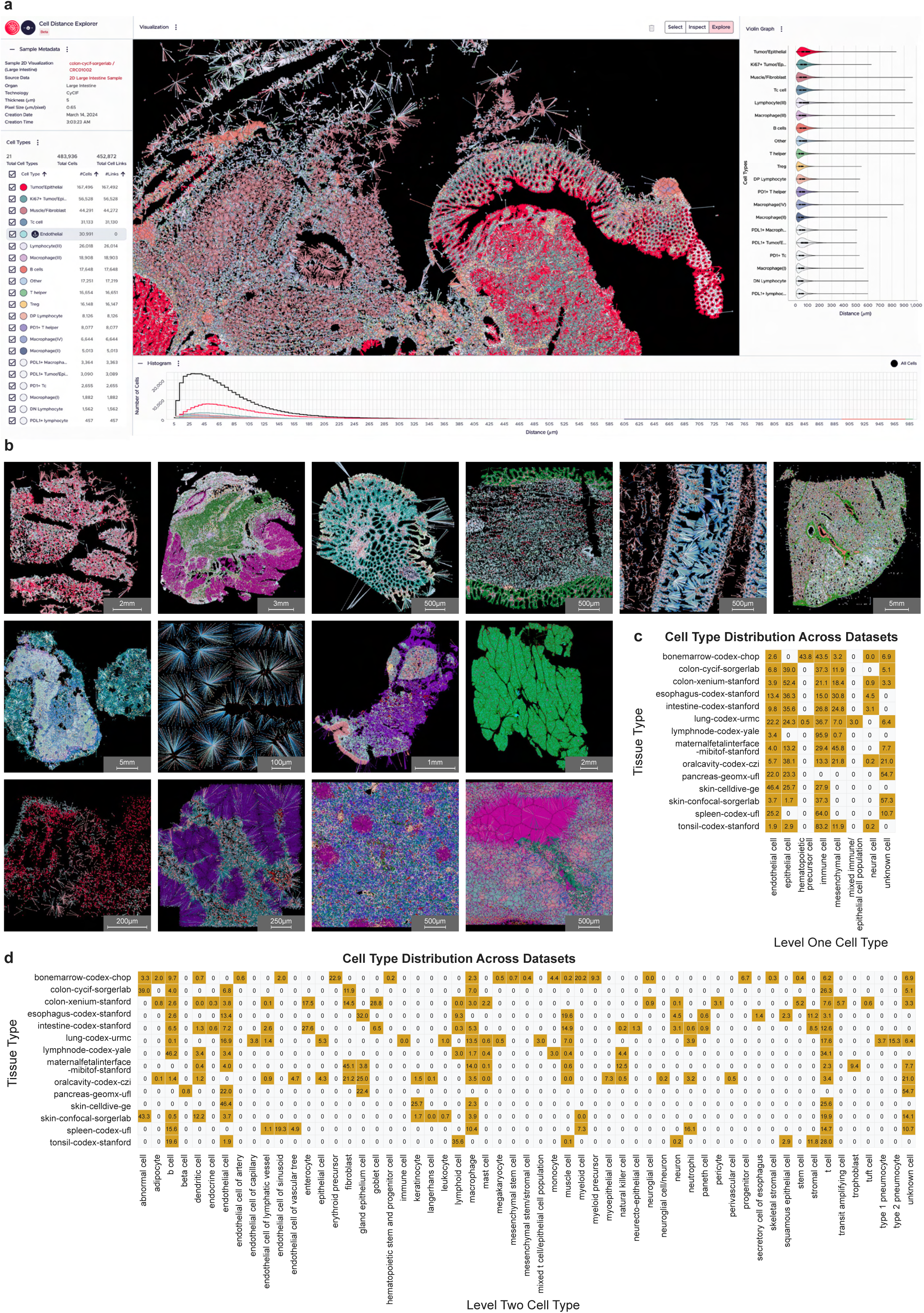
Cell Distance Explorer and overview of collected studies and datasets. **a.** A colon dataset visualized in the online tool, Cell Distance Explorer, showing cell-to-nearest-endothelial-cell linkages in the central panel, and distance distributions per cell type visualized as a violin plot (right) and histogram (bottom). **b.** A sample dataset from each collected study is visualized in the Cell Distance Explorer. Top row (left to right): bonemarrow-codex-chop, colon-cycif-sorgerlab, colon-xenium-stanford, esophagus-codex-stanford, intestine-codex-stanford, lung-codex-urmc. Middle row (left to right): lymphnode-codex-yale, maternalfetalinterface-mibitof-stanford, oralcavity-codex-czi, pancreas-geomx-ufl. Bottom row (left to right): skin-celldive-ge, skin-confocal-sorgerlab, spleen-codex-ufl, tonsil-codex-stanford. **c.** Binary heatmap showing presence of cell types at level L1 across collected studies. Numbers show the percentage of the cell type within that collected study. **d.** Binary heatmap showing presence of cell types at level L2 across collected studies. Numbers show the percentage of the cell type within that collected study.

## Results

### Cell Distance Explorer, a web-based interactive tool for spatial cell graph exploration

We developed a web-based interactive tool, Cell Distance Explorer (CDE, https://humanatlas.io/user-story/5), see **Fig. 1a** for a screenshot of the user interface with a sample colon dataset. The CDE allows researchers to compute cell-to-nearest-anchor-cell distance distributions for their cell types of interest and visualize them in an interactive environment. As part of development, we demonstrated the tool to several researchers in the field of computational biology and bioinformatics, and their responses and feedback helped prioritize and optimize tool functionality.

CDE users upload a dataset containing x, y, z coordinates of cells and their assigned cell type labels as a CSV file; z is needed if 3D data exists but optional. Additionally, users can define several parameters such as maximum distance threshold and pixel size of their dataset that are used for computation and visualization. Users can change colors of assigned cell types and delete irrelevant cells in edit mode. They can download the final visualizations in PNG/SVG format and data as cells.csv, edges.csv, and color map.csv files for further investigation.

In addition to the web-based tool, a Python package is available so users can use the tool in Jupyter notebooks, see **Code Availability**. Moreover, the CDE is available as a lightweight web component which can be embedded in HTML webpages, see **Code Availability**. Note that the tool can also be used to compute the distance of cells to other entities for which spatial coordinates and labels are available (e.g., locations of named subcellular biomarkers or the outline of labelled functional tissue units).

### Collecting spatially resolved omics datasets containing endothelial cells

We compiled a collection of already published spatially resolved omics datasets from various atlassing efforts for a meta study. The resulting data collection comprises 399 spatial transcriptomics and spatial proteomics single-cell datasets from 14 different studies (see **Methods**) comprising 12 different tissue types and seven spatial omics technologies (CODEX, Cell Dive, 3D CyCIF by serial-section reconstruction, 3D CyCIF by confocal microscopy, Xenium, MIBI-TOF, GeoMx). The complete data collection contains 47,349,496 cells with spatial 2D or 3D coordinates (cell centroids) and assigned cell type labels. All 295 original cell type labels were harmonized into a 3-level typology and crosswalked to Cell Ontology. Complete listing of collected studies, including number of datasets, cells, and cell types per study, is presented in **Table 1**. A sample dataset, visualized in the Cell Distance Explorer, from each study is shown in **Fig. 1b**.

**Table 1.**
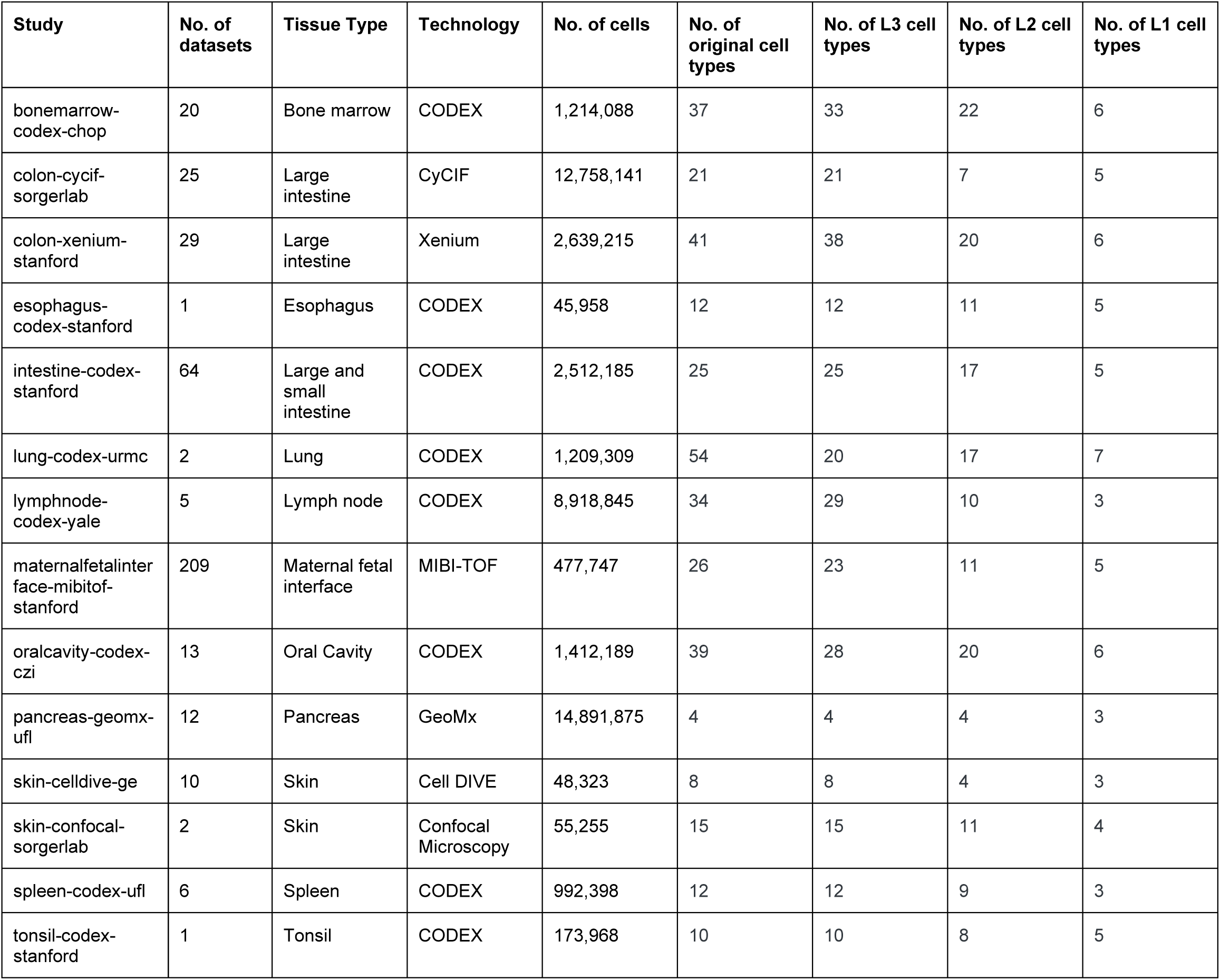
Listing of all collected studies, relevant tissue and technology information, total number of cells and datasets per study, and number of unique cell types present in the datasets across different cell type levels.

The spatial size, location, and orientation of tissue specimens were manually registered using the Registration User Interface^16^ in collaboration with organ-expert data providers and were published in the Exploration User Interface^16^. A total of 139 datasets, corresponding to 107 tissue blocks from 7 tissue types, 38 donors, and 8 data providers were registered (see **Supplementary Table 1)**. The registered collection can be explored in the Exploration User Interface at https://hubmapconsortium.github.io/hra-registrations/vccf-federated.

### Harmonizing cell types across the data collection

The initial cell type labels assigned by the original authors of different datasets contained several issues such as differences in spelling and capitalization, presence of different cell type names for the same cell type, and usage of cell type labels with different levels of specificity in the typology. As the first step for cleaning the collected data, we harmonized the cell type labels across all 399 datasets: 1) We converted all labels to lower capitalization and used a consistent format across all cell types, 2) We used a 3-level hierarchical cell type typology in terms of specificity of labels (Level Three Cell Type (L3), Level Two Cell Type (L2), and Level One Cell Type (L1)) 3) We mapped each cell type label at all three levels of typology to Cell Ontology^18,19^.

L3 is the most specific cell type categorization possible, given the omics technology and the antibody panel used in the study, while L1 is the most general label (categorizing all cell types into 8 categories: *endothelial*, *epithelial*, *hematopoietic precursor*, *immune*, *mesenchymal*, *neural*, *mixed immune/epithelial*, and *unknown*). The L1 categories were initially defined to classify all cell types as *endothelial*, *immune*, *epithelial*, *mesenchymal*, and *unknown*, but other three categories were added due to the presence of such cell types in some studies. **Fig. 1c** shows the presence of cell types at L1 across studies. Since the cell type categories are inconsistent across studies, the three-level cell type typology is defined to enable cross-study comparisons, for example, L2 classification can help comparing *B cells* and *T cells* across studies whereas L3 classification can enable comparing *immune cells* and *epithelial cells* across studies. **Fig. 1d** shows the presence of cell types at L2 across studies.

The original cell type labels consisted of 295 unique cell type categories (without any data cleaning), whereas the harmonized dataset has 162 unique labels for L3, 57 unique labels for L2, and 8 unique labels for L1. At L1 classification, the most shared cell type categories across studies are *endothelial*, *epithelial*, *immune*, and *mesenchymal*. At L2 classification, *unknown cell*, *T cell*, *B cell*, *abnormal cell*, and *endothelial cell* are the five most shared cell types across the studies. It should be noted that 8,147,411 cells in the *unknown cell* category are from the pancreas-geomx-ufl datasets; the total number of *unknown cells* at L2 classification is approximately 9.4 million across the entire collection. The most abundant cell types across the collection at L1 classification are (in order): *immune*, *epithelial*, *unknown*, *endothelial*, and *mesenchymal*. For *ECs*, all cells are categorized as *endothelial cell* at L1 classification, but there are other *EC* categories available for L2 and L3 classification: *endothelial cell*, *endothelial cell of artery*, *endothelial cell of capillary*, *endothelial cell of lymphatic vessel*, *endothelial cell of sinusoid*, and *endothelial cell of vascular tree*. The complete listing of all cell types, including original cell type labels and harmonized cell type labels, is available at https://github.com/cns-iu/hra-cell-distance-analysis/blob/main/data/mapping_files/generated_cell_type_complete_crosswalk.csv.

### Distance analysis across unique regions in a tissue type

To characterize the cell-to-nearest-endothelial-cell distance distributions across different regions within a tissue, we looked at three studies where specific unique region information was available for the datasets: intestine-codex-stanford, pancreas-geomx-ufl, and oralcavity-codex-czi studies. For datasets in each study, an appropriate *endothelial cell* category is defined at each classification level as the anchor cell type and euclidean distance is computed for each cell to the nearest endothelial cell, which defines the linkage (or edge) between the cell and the nearest *endothelial cell*.

Studying endothelial cell-neighbor cell distances at a population level can reveal a number of biological phenomena. In general, a small interquartile range (IQR) spread of endothelial cell-neighbor distances suggests that such cell types may have a more predictable and uniform positioning relative to blood vessels or there exist greater consistency across donors. Such uniform cell organization of specific cell types may suggest a tight biological regulation of cell placement due to specific functional needs of such cell types related to their nearness to blood vessels. Conversely, a larger IQR spread suggests a more variable spatial organization of cells, based on how such cell types can be found at many different distances from the vessels. Such variable cell organization may suggest a more flexible tissue architecture, or multiple functional needs of such cell populations that go beyond their dependence on blood vessel proximity, or greater donor-donor variations. Additionally, lower median distances to endothelial cells generally suggest such cells have higher metabolic demands and require immediate access to blood vessels, potentially for faster and easier circulation, rapid surveillance, and nutrient access. Conversely, cells with higher median distances may suggest vascular-independent functional priorities or structural use instead of a functional use. Furthermore, more complex tissue architecture can also lead to higher variable distances.

For the intestine-codex-stanford dataset, the 64 datasets are taken from eight unique regions in the small and large intestines: Ascending, Descending, Duodenum, Ileum, Mid Jejunum, Proximal Jejunum, Transverse, Sigmoid (**Fig. 2a**). For pancreas-geomx-ufl, 12 datasets are taken from four unique regions: Head, Neck, Body, Tail (**Fig. 2b**). For oralcavity-codex-czi, the 13 datasets are taken from five unique regions in the oral cavity (mouth): Buccal Mucosa and Minor Salivary Glands, Gingiva, Parotid, Submandibular, Tongue (**Fig. 2c**).

**Figure 2.**
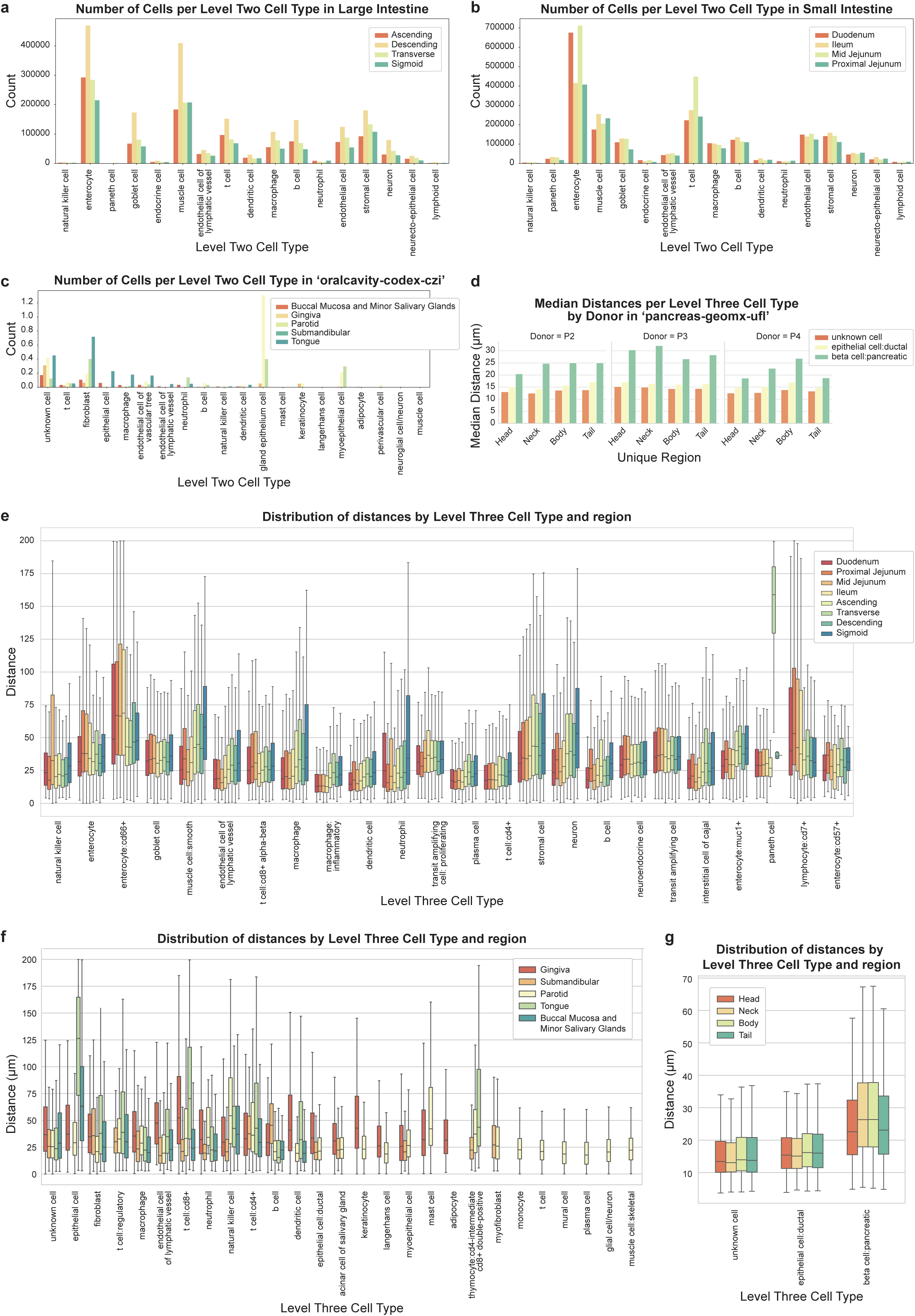
Distance analysis across unique regions in a tissue type. **a.** Bar graph depicting number of cells per L2 cell type for each unique region in large intestine in intestine-codex-stanford datasets. **b.** Bar graph depicting number of cells per L2 cell type for each unique region in small intestine in intestine-codex-stanford datasets. **c.** Bar graph depicting number of cells per L2 cell type for each unique region in oral cavity in oralcavity-codex-czi datasets. **d.** Bar graph depicting median distances per each unique region in pancreas, faceted by donors in pancreas-geomx-ufl datasets. **e.** Boxplot depicting cell-to-nearest-endothelial-cell distance distributions per L3 cell type for each unique region in small and large intestine in intestine-codex-stanford datasets. **f.** Boxplot depicting cell-to-nearest-endothelial-cell distance distributions per L3 cell type for each unique region in the oral cavity in oralcavity-codex-czi datasets. **g.** Boxplot depicting cell-to-nearest-endothelial-cell distance distributions per L3 cell type for each unique region in the pancreas in pancreas-geomx-ufl datasets.

**Intestine-codex-stanford**^17,20^: At the highest level of resolution (L3) we observed that *inflammatory macrophage* has the lowest median distance for both small and large intestines (**Fig. 2e**). This is particularly interesting because of our prior finding that *inflammatory macrophages* are restricted to the mucosa and further enriched towards the top of the crypt^17,21^. Our endothelial-specific finding can be interpreted based on this initial finding for two reasons. First, the top of the intestinal crypt is also an area of dense capillaries^22^. Second, because the *inflammatory macrophages* are primarily enriched in this area of the mucosa, the *endothelial-inflammatory macrophage* distance will be lower than tissue average where its non-inflammatory *macrophage* cell type falls in line as this is found across different tissue compartments (**Fig. 2e**).

In addition to the decreased median distance, *inflammatory macrophages* had a smaller IQR as compared to other cell types such as the *macrophage* cell type (**Fig. 2e**). This change in variance can be understood as since *inflammatory macrophages* are restricted to a certain area of the intestine, their distance to endothelial cells will be more constant. Consequently, there are different endothelial cell densities across the intestine with the highest density occurring in the villus. Similar cell types that have highly conserved spatial locations in the intestine and not found across the intestine are *plasma cells* and *dendritic* cells that also have a lower IQR than other more spatially ubiquitous cell types. This indicates that cell types that are restricted to certain environments will have smaller IQR than others distributed across various environments.

Another global observation that we can make is that the median distance for many cell types is lower within the small intestinal regions and compared to the large intestine regions (**Fig. 2e**). This is consistent with our prior findings that suggests lower endothelial cell percentages in the large intestine as compared to the small intestine^17^. It is also consistent with the function of the small intestine that is involved in nutrient extraction. While we only highlighted a few of our observations, we can observe significant biology of organs from calculating and evaluating endothelial cell-cell distances.

**Pancreas-geomx-ufl**^23–34^: At L3 resolution, we observed that *pancreatic beta cells* have a higher median distance and IQR than other cell types, likely impacted by their known secretion of insulin into the vasculature and the presence of *beta cells* within islets (> 8 *beta cells*), clusters (2-8 *beta cells*) and single *beta cells* randomly scattered throughout the parenchyma (**Fig. 2g**). We observed that the neck and body regions have a higher median distance and IQR than the head and tail regions when considering all datasets combined. At the individual donor level, this pattern holds true for P4 but not for P2 and P3 (**Fig. 2d**). There is no apparent biomedical explanation given the morphological similarities of islet density between body and tail regions while studies of islet density in the human pancreas neck region have not been reported.

**Oralcavity-codex-czi**^35–37^: *Mucosa:* The tongue epithelium is thick, keratinized, and stratified, especially on the dorsal surface, requiring a deeper vascular network compared to glandular tissue. This architecture drives an increased median distance and IQR between *ECs* and *epithelial* or *immune* cells. The elevated *EC* distance for *CD8+ T cells* at L3 resolution likely reflects *intraepithelial lymphocytes (IELs)*, a specialized *CD8+* subset that resides within the epithelium and therefore farther from vessels (**Fig. 2f**). Gingiva, similarly keratinized but under constant microbial and mechanical stress, is enriched in fibrovascular tissue beneath its dense epithelial barrier^35,36^. The median distance and IQR for *keratinocytes* is much higher in gingiva relative to glands, reflecting a more variable spatial organization of these cells in gingiva compared to the parotid.

*Glands:* Unlike the layered, keratinized tongue epithelium, the glandular epithelium is simple and less stratified, allowing *IELs* (*CD8+ T cells* within *epithelial* cell types) to remain in closer proximity to *ECs*, reflected in their lower median distance and IQR compared to tongue *IELs*. For example, the parotid gland is a densely innervated serous secretory organ, with neurovascular bundles serving acini and ducts. This is reflected in the abundance of *acinar cells* and their close *EC* proximity. *Neural cell* proximity to *ECs* likely reflects parasympathetic inputs. The short *B cell*–*EC* distance may capture *gland-resident plasma cells* supporting IgA secretion (**Fig. 2f**). The submandibular gland, with mixed serous–mucous output, is known to harbor *IELs* positioned near ductal epithelium and vasculature also as expected.

The variation in *CD8+ T cell* distances from *endothelial cells* across oral tissues likely reflects differences in epithelial architecture and stromal organization, which is well known but not well-studied in this context. In mixed sites such as buccal mucosa which contains thick, stratified epithelial and minor salivary glands in the stroma, the distributions likely represent an averaging of longer-range *IEL* positions in epithelium with shorter-range *resident T cell* positions in lamina propria or periductal compartments. Together, these spatial differences suggest that local tissue architecture constrains the positioning of *T cells* relative to vasculature, with implications for how immune surveillance is organized across oral niches^38^.

### Distance analysis across disease conditions in same tissue type

To characterize the changes in the cell-to-nearest-endothelial-cell distance distributions across different disease conditions in a tissue, we looked at five studies where specific disease condition information was available for the datasets: lung-codex-urmc, skin-confocal-sorgerlab, bonemarrow-codex-chop, colon-xenium-stanford, and colon-cycif-sorgerlab studies. For datasets in each study, an appropriate *endothelial cell* category is defined at each classification level as the anchor cell type and euclidean distance is computed for each cell to the nearest *endothelial cell*, which defines the linkage (or edge) between the cell and the nearest *endothelial cell*.

**Lung-codex-urmc**^6^: In prior work^6^, we looked at similar regions of interest (bronchiole and an accompanying small pulmonary artery) in a lung tissue from a healthy adult and a lung tissue from a child with bronchopulmonary dysplasia (a chronic lung disease following premature birth). In brief, we found increased cellularity with a predominance of *CD4+ lymphocytes* and *myeloid immune cells* positioned in close proximity to lung vasculature in the diseased lung.

Here, we analyzed the same samples with improved cell segmentation and more granular cell type annotation. At the lowest resolution (L1), we observed that *immune cells* had a higher median distance and IQR in the disease sample compared to the normal sample, the increased vascular distances and variability likely reflecting pathological changes leading to vessel remodeling (**Fig. 3a**). We observed *muscle cells* have a closer vascular proximity in disease tissue. Conversely, *macrophages* have a higher median distance and IQR in disease tissue, likely reflecting infiltration into inflamed interstitium (**Fig. 3b**). At the highest resolution (L3), we observed that *T cell* positioning shifted between conditions: *CD4+ alpha beta T cells* were more distant in normal tissue whereas *CD8+ T cells* were more distant in disease tissue (**Fig. 3c**).

**Figure 3.**
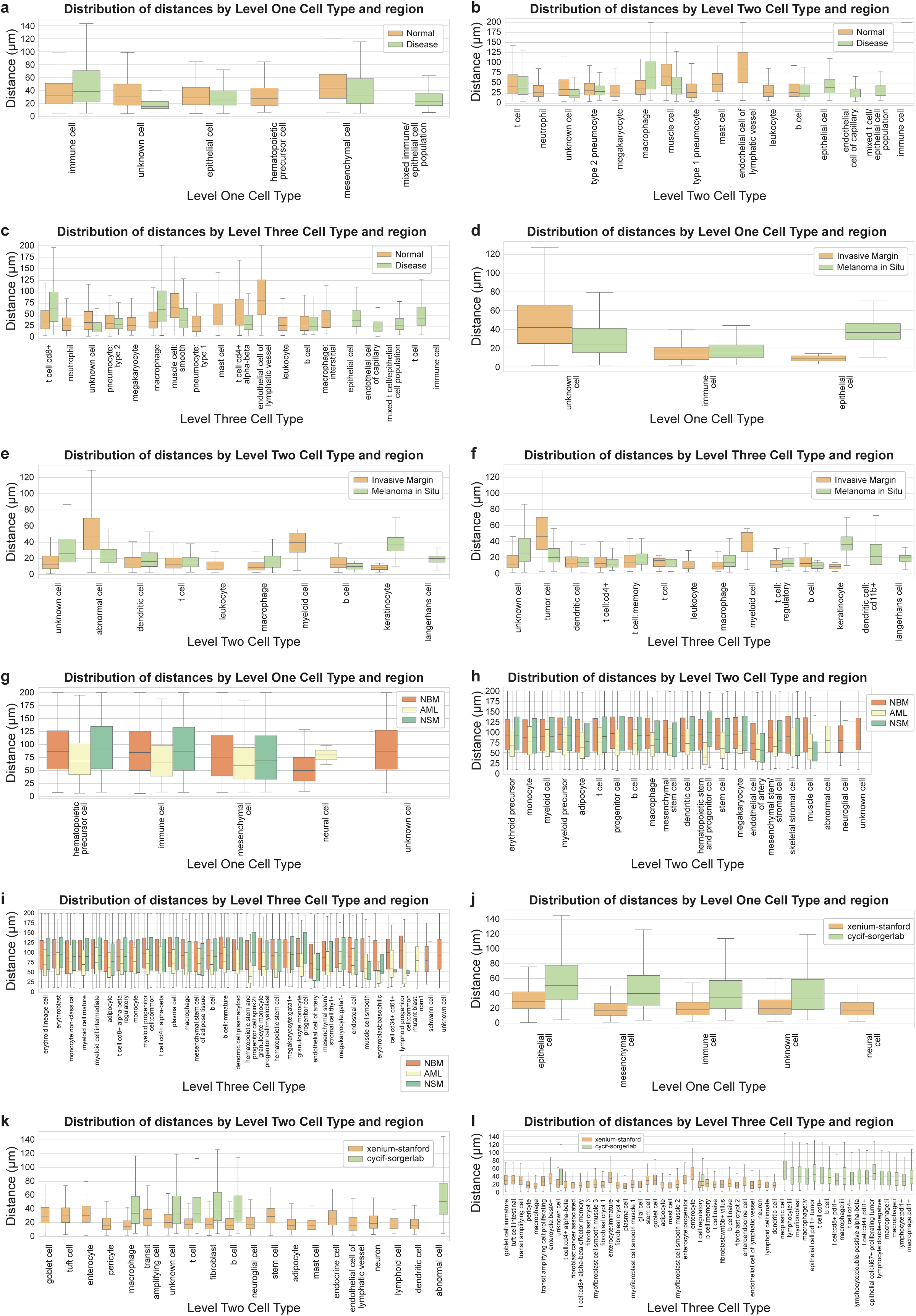
Distance analysis across disease conditions in the same tissue type. **a-c.** Boxplots depicting cell-to-nearest-endothelial-cell distance distributions per cell type for each cell type level for normal and disease region in lung in lung-codex-urmc datasets. **d-f.** Boxplots depicting cell-to-nearest-endothelial-cell distance distributions per cell type for each cell type level for invasive margin and melanoma in situ region in skin in skin-confocal-sorgerlab datasets. **g-i.** Boxplots depicting cell-to-nearest-endothelial-cell distance distributions per cell type for each cell type level for NBM, NSM, and AML regions in bone marrow in bonemarrow-codex-chop datasets. **j-l.** Boxplots depicting cell-to-nearest-endothelial-cell distance distributions per cell type for each cell type level for Xenium and CyCIF regions in colon in colon-xenium-stanford and colon-cycif-sorgerlab datasets.

Abnormal immune activity causes vascular leakiness and edema, further thickening membranes while mucus and debris accumulation creates mismatch between flow of air and blood; persistent inflammation ultimately leads to fibrosis. In bronchopulmonary dysplasia, extravascular immune cell aggregates thicken gas exchange membranes, creating compressed yet variable distance distributions to *endothelial cells*.

**Skin-confocal-sorgerlab**^39^: This dataset is unique by featuring multi-layered full volume 3D representations of cells and blood vessels acquired over 170 z-planes by high resolution confocal microscopy, so all distances are computed in a 3D space.

We noticed that *T cells* and *B cells*–which are continuously undergoing recruitment from the circulatory system–have a relatively low median distance in both regions (**Fig. 3e**). Additionally, the *epithelial cells* have a higher median distance and IQR in melanoma in situ compared to invasive margin potentially indicating a more variable spread of *epithelial cells* in melanoma in situ (**Fig. 3d**). The majority of *immune cells* in the invasive margin are *resident T cells* (compared to just half in the melanoma in situ). Upon reviewing the primary data, we observed some instances of *resident T cells* migrating large distances away from *endothelial cells* and invading deep into the tumor. We observed that there is a higher *immune* and *abnormal/tumor cell* population in the invasive margin dataset relative to melanoma in situ dataset. This is consistent with the fact that the invasive margin is a more progressed level of disease whereby we expect tumor proliferation into the dermis leading to recruitment of *immune cells* followed by an immune response. For *abnormal* (*tumor cells*), we saw an increase in median distance with disease progression (melanoma in situ: 20.3 µm vs 46.41µm invasive margin; **Fig. 3f**). In melanoma in situ, transformed *melanocytes* (denoted as ‘*tumor*’ due to MART1 positive staining) were characteristically observed to be arranged as a single layer along the dermal epidermal junction and have close proximity to the dense vascular system just below in the dermis. In contrast to the melanoma in situ, the invasive margin consisted of *tumor cells* forming large colonies whereby centrally-located *tumor cells* are positioned deep behind many cell-distances away from the nearest endothelial vessel, thus, resulting in a larger median distance. We also observed a big difference in the *keratinocyte* median distance in the two samples although this is potentially due to the marked decrease in *keratinocytes* in the invasive margin (**Fig. 3f**).

**Bonemarrow-codex-chop**^40^: We observed that the AML samples showed consistently lower median distances across all major cell types (*immune cells*, *mesenchymal cells*, and *hematopoietic precursor cells*) compared to NBM and NSM samples (**Fig. 3g**). This spatial compression of cells likely reflects the effect of leukemic infiltration disrupting the normal bone marrow architecture. The similarity between the NBM and NSM distributions across all cell types likely indicates that systemic lymphoma doesn’t necessarily alter spatial organization with respect to *sinusoidal endothelial cells* (**Fig. 3h**). Notably, the *smooth muscle cells* showed closer vascular proximity in NBM samples compared to both disease states, suggesting a spatial expansion of vasculature in disease states (**Fig. 3i**).

**Colon-xenium-stanford**^41^ **and colon-cycif-sorgerlab**^42^: We also compared colon samples from two different studies. The Xenium sample contains 29 serial-sections from a colon polyp, whereas the CyCIF sample contains 25 serial-sections spaced 25 µm apart from a colorectal cancer sample. We considered each section as a separate non-aligned tissue and computed distances in a 2D system.

In general, at the lowest resolution, the Xenium dataset has a lower and tighter distance distribution than the CyCIF dataset (**Fig. 3j**). Similarly, for major cell type categories (*B cell*, *T cell*, *fibroblast*, and *macrophages*), we observed a roughly 3-fold increase in median distances for the CyCIF dataset as compared to the Xenium dataset (**Fig. 3k**). At the highest resolution (L3), we observed that the median distance is similarly high for *neoplastic cells*, *PDL1+ tumor associated epithelial cells*, and *Ki67+ proliferating tumor epithelial cells* in the CyCIF dataset (**Fig. 3l**). In the Xenium dataset, median distance for *cancer associated fibroblasts* is 15.4 µm which is fairly close to other *fibroblast* types in the dataset; *plasma cells* have the lowest median distance of 13.9 µm whereas the largest median distance is of different types of *enterocytes*, *goblet cells*, *transit amplifying cells*, and *intestinal tuft cells*. For the CyCIF dataset, different types of *lymphocytes* and *macrophages* have a varying median distance lying in the range of approximately 42-55 µm (**Fig. 3l**).

However, these differences were not statistically significant between the two imaging modalities despite the 2-3-fold differences in median distances. We also note that the two datasets, although coming from the same organ type, are from two entirely different donors with different severities of disease. For example, the Xenium dataset was from a colon polyp–usually taken at early signs of cancer–whereas the CyCIF dataset was from *stage IIIb* adenocarcinoma cancer. Thus, the above analysis may demonstrate differences in cell distributions across disease severities. Furthermore, the Xenium region size is several orders of magnitude smaller whereas CyCIF sampled a larger and a potentially more representative area of tissue.

### Distance analysis across individual datasets in other tissue types

To further characterize the cell-to-nearest-endothelial-cell distance distributions in different tissues, we looked at six studies: lymphnode-codex-yale, spleen-codex-ufl, tonsil-codex-stanford, esophagus-codex-stanford, skin-celldive-ge, and maternalfetalinterface-mibitof-stanford studies. For datasets in each study, an appropriate *endothelial cell* category is defined at each classification level as the anchor cell type and euclidean distance is computed for each cell to the nearest endothelial cell, which defines the linkage (or edge) between the cell and the nearest *endothelial cell*.

**Lymphnode-codex-yale**^43–48^: We observed substantial inter-donor variability, with the median distances across all cell types exceeding 55 µm in the LN00560 donor sample versus below 50 µm in the LN27766 donor sample. Distances are more varied in the other three samples, for example, in the LN22921 sample, the *muscle cells* have a median distance of 28.4 µm and the *B cells* that of 72.3 µm (**Fig. 4a**). *Mast cells* in the LN00560 sample have a relatively very high median distance of 96.2 µm, likely explained by the function and density of these respective cells, i.e. *smooth muscle cells* compose the wall of vasculature whereas mast cells are fewer (scattered) compared to *lymphocytes* in a lymph node.

**Figure 4.**
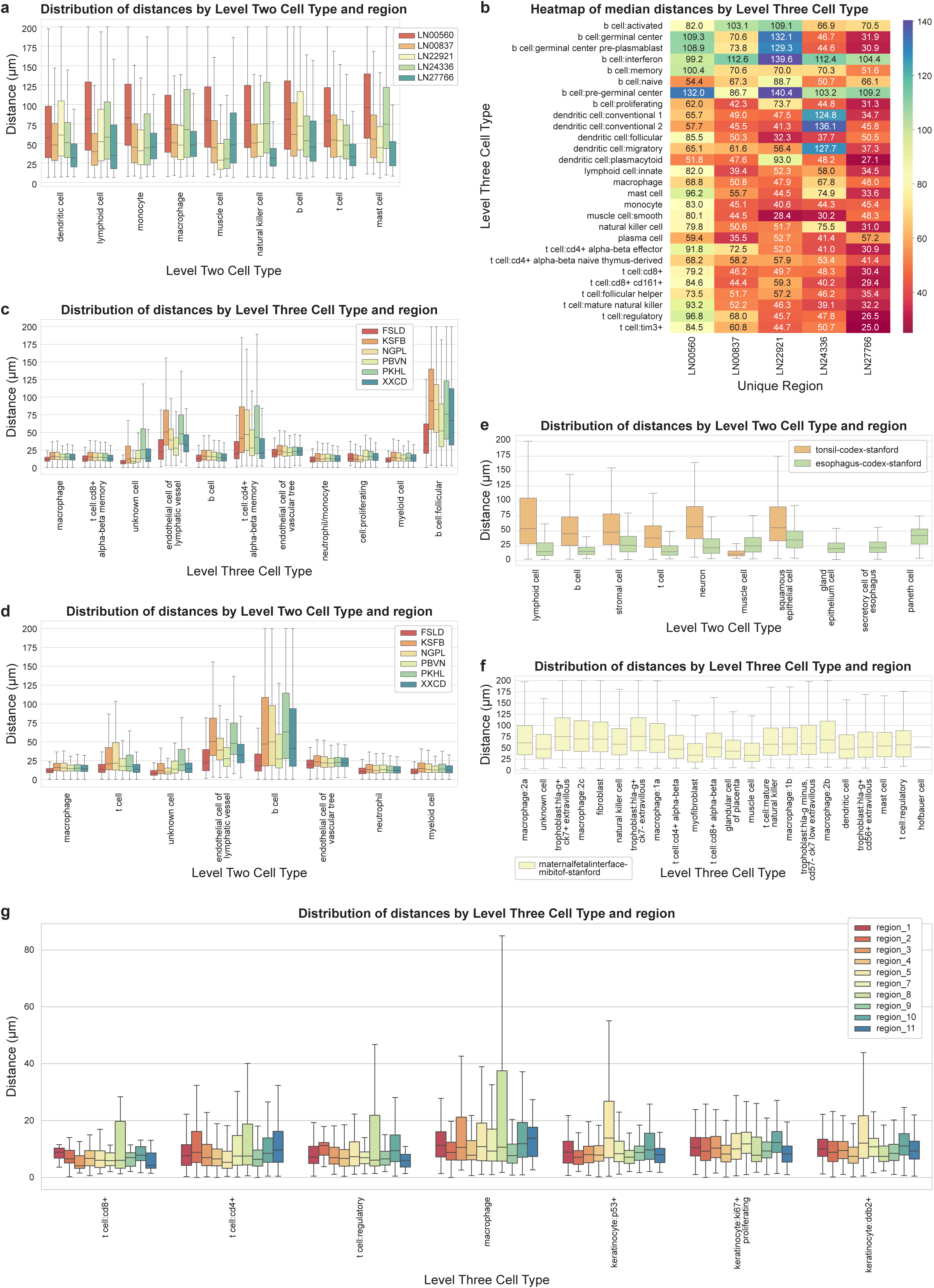
Distance analysis across individual datasets in other tissue types. **a.** Boxplots depicting cell-to-nearest-endothelial-cell distance distributions per L2 cell type for different lymph node regions in lymphnode-codex-yale datasets. **b.** Heatmap showing median distances for L3 cell types for lymph node data in lymphnode-codex-yale datasets. **c.** Boxplots depicting cell-to-nearest-endothelial-cell distance distributions per L3 cell type for different spleen regions in spleen-codex-ufl datasets. **d.** Boxplots depicting cell-to-nearest-endothelial-cell distance distributions per L2 cell type for different spleen regions in spleen-codex-ufl datasets. **e.** Boxplots depicting cell-to-nearest-endothelial-cell distance distributions per L2 cell type for esophagus and tonsil regions in esophagus-codex-stanford and tonsil-codex-stanford datasets. **f.** Boxplots depicting cell-to-nearest-endothelial-cell distance distributions per L3 cell type in maternalfetalinterface-mibitof-stanford datasets. **g.** Boxplots depicting cell-to-nearest-endothelial-cell distance distributions per L3 cell type for different skin regions in skin-celldive-ge datasets.

At the highest resolution (L3), different types of *T cells* have a relatively lower median distance than different types of *B cells*–indicating consistently closer vascular proximity in *T cell* subtypes compared to *B cell* subtypes (**Fig. 4b**). Some B cell subtypes like the *germinal center related B cells* showed a high median distance across donors, possibly due to their positioning in follicle centers. Additionally, some rarer *B cell* subtypes like *pre-germinal center B cell* and *interferon B cells* also showed high distance distributions, but the low cell counts might affect statistical reliability. *T cells* are known to predominate over *B cells* in the paracortex, which is also rich in high endothelial venules. *B cells* are more centralized in primary and secondary follicles. Compared to the middle-aged donors, the lymph nodes from the most elderly donor showing greater distance of various cell types likely reflect lower overall cellularity due to aging. It’s unclear why this is also seen in the youngest donor, but may be donor specific. Since there is only one donor younger than 30 years old, it is difficult to extrapolate further from the dataset.

**Spleen-codex-ufl**^49,50^: We observed that most splenic cell populations localized within 50 µm of *sinusoidal endothelial cells*, likely reflecting the highly vascular architecture of spleen required for blood filtration. Notably, an exception to this pattern was *B cells*, particularly *follicular B cells*, which showed consistently higher median distance and IQR across samples, possibly due to their organization in white pulp follicles distant from the red pulp vasculature (**Fig. 4c**). *T cells* demonstrated closer vascular proximity compared to *B cells*, similar to lymph node dataset (**Fig. 4d**). Among *T cell* subtypes, *CD8+ alpha beta memory T cells* showed lower vascular positioning compared to *CD4+ alpha beta memory T cells* across all six samples, potentially due to different microenvironmental niches within splenic compartments.

**Tonsil-codex-stanford and esophagus-codex-stanford**^51,52^: We analyzed the tonsil and esophagus datasets together that come from the same study^51^, each containing a single dataset. The tonsil dataset contains a much higher *lymphoid cell*, *T cell*, and *B cell* population compared to the esophagus dataset, characteristic of secondary lymphoid organs, while the esophagus dataset contains a higher *endothelial cell* population, possibly due to barrier tissues requiring extensive vascularization for nutrient supply and immune surveillance.

We observed that the esophagus dataset has a lower median distance, IQR, and outlier range across all cell types compared to the tonsil dataset, likely reflecting the compact *epithelial*-*stromal* architecture of esophagus where cells might maintain close vascular proximity for efficient nutrient exchange and immune monitoring of the mucosal barrier (**Fig. 4e**). We also observed that the median *T cell* distance is much lower than *B cell* in tonsil, while it is much closer to each other in the esophagus. This might reflect a difference in the functional specialization of *T cells* in tonsil, which is a secondary lymphoid organ requiring rapid immune response, versus esophagus which is a barrier tissue requiring immune surveillance and nutrient supply. It should be noted that since both datasets include a single sample, the conclusions might lack statistical reliability which would require more samples.

**Skin-celldive-ge**^53^: In prior work^53^, we found significant differences in the average distance of the nearest *endothelial cell* to *immune cells*, with distances in 3D half that found in 2D (∼56 µm vs 108 µm on average). Prior work in breast skin tissue also demonstrated, using confocal imaging^54^, that *T cells* form perivascular sheaths throughout the dermis and reside within 15 µm distance from *endothelial cells*. We observed that most *T cells* were located within 20 µm of nearest *endothelial cells* across all regions (**Fig. 4g**). We also observed a much higher IQR for *macrophages* and different *T cell* subtypes in region 8 compared to other regions likely due to reduced number of overall inflammatory cells in this donor skin compared to others; notably, this sample comes from a donor with HIV infection that is associated with reduction in *T cells*. We also noticed a higher IQR for *P53+* and *DDB2+ epithelial cells* in region 5 relative to other regions; this sample comes from the youngest donor (33) across all regions, reflecting well-preserved epidermal and dermal architecture and suggesting higher regenerative capacity relative to older donors. Region 5 and region 8 also have a higher coefficient of variability (100.2% and 106.3%) whereas region 1 has the lowest (54.9%) – region 5 is the youngest donor, region 8 is a donor with HIV, and region 1 is a scalp sample from the oldest donor (72) with rheumatoid arthritis.

**Maternalfetalinterface-mibitof-stanford**^55^: We observed that *trophoblasts*, which play a key role in early pregnancy by facilitating embryo implantation and form the majority of the placenta, had the highest median distance compared to other cell types, particularly driven by *hla-g+ ck7+ extravillous* and *hla-g+ ck7-extravillous trophoblasts* (**Fig. 4f**). This may be due to the deep invasion of these cells in the maternal uterine lining to anchor the placenta, remodel the maternal vasculature, and ensure adequate blood and nutrient supply to the fetus. Conversely, the *muscle cells* showed very close vascular proximity, likely due to their structural association with vessel walls and requirement for direct vascular support. Similarly, *myofibroblasts* also showed very close vascular proximity, likely providing high tissue repair and generation support.

### Endothelial cell-focused neighborhood analysis

While single-cell cell-cell interaction analysis provides critical insights, these analyses often overlook the local spatial context in which each cell resides, potentially altering our interpretation of cellular behavior. This is particularly relevant for endothelial cells, as vasculature spans multiple tissue regions and subregions. In the intestine, for example, vasculature is present across the mucosa, submucosa, and muscularis externa. Within each layer, distinct *EC*-centric neighborhoods exist, each composed of specialized cellular communities that support the unique functional demands of their microenvironment.

In normal cell-cell interaction analysis, these are lumped together which diminishes unique cell-cell interactions by averaging them out. In this context, spatial organization is critical for understanding how vascular structures behave and contribute to tissue-specific functions. To capture the nuances between EC “neighborhoods,” we investigated the vascular organization across multiple spatial scales by examining how the composition of neighboring cells varies around *ECs* as well as the co-localization of these cellular neighborhoods. Identifying and characterizing these cellular neighborhoods enhances our ability to define the vasculature as an integrated network of functional units within tissue subregions.

We have previously developed an algorithm to establish cellular neighborhoods based on one cell type and identified different *T cell* environments within a tumor^56^. To identify vasculature-based neighborhoods, we used this cell-centric neighborhood technique on the intestine-codex-stanford data^17^ (**Fig. 5a**). Specifically, we identified the 50 nearest neighbors for each *EC* (**Fig. 5b**) and clustered these local microenvironments using K-means clustering, resulting in 11 distinct *endothelial cell*-centric neighborhoods (**Fig. 5c-e**).

**Figure 5.**
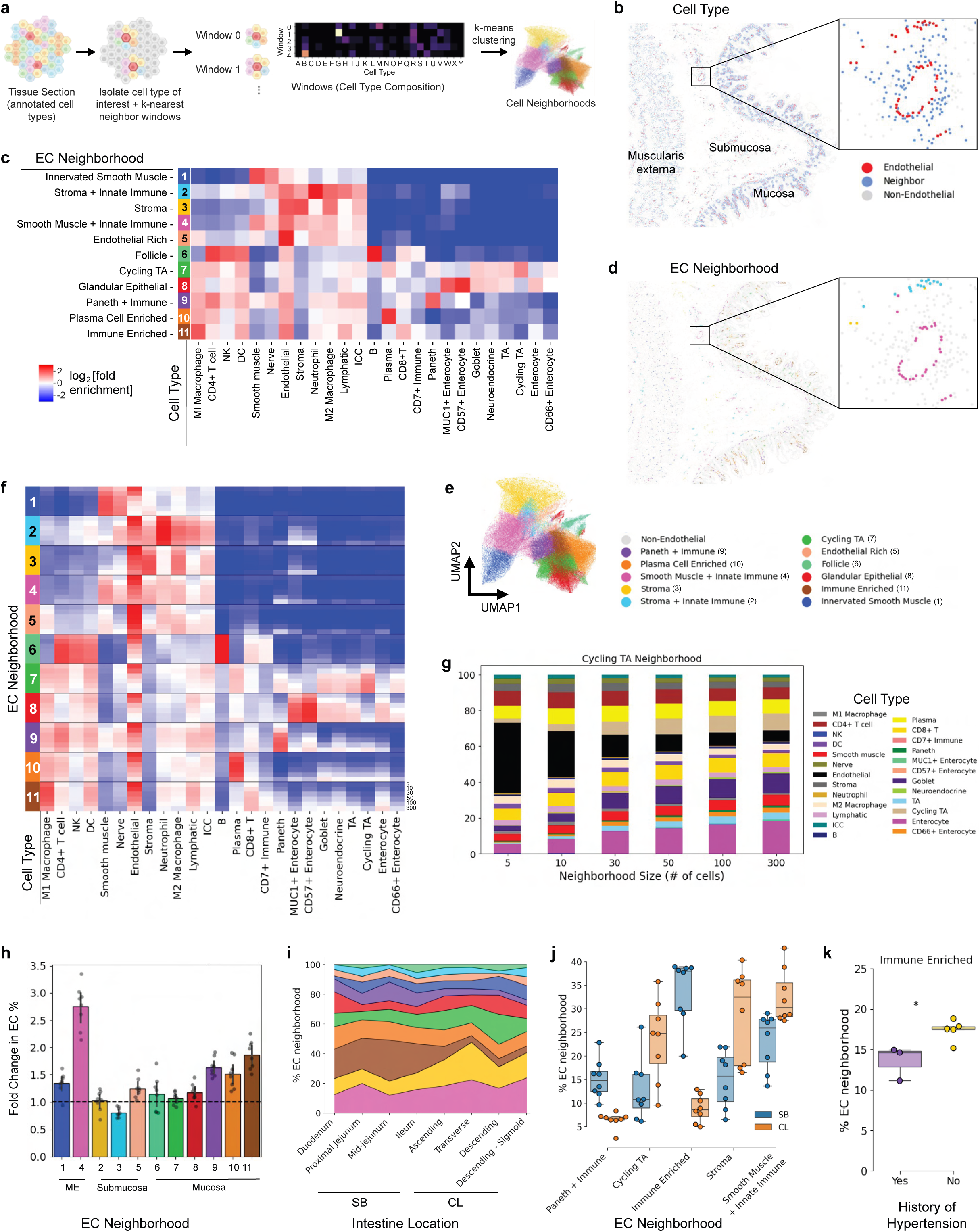
Single-cell neighborhood analysis of endothelial cells (ECs) in the healthy human small intestine and large intestine. **a.** Schematic of cell type-centric neighborhood analysis pipeline. **b.** Representative image of the intestine highlighting ECs and their 50 nearest cell neighbors. **c.** Heatmap depicting cell type abundance within each neighborhood. **d.** ECs with neighborhood labels mapped to tissue coordinates. **e.** UMAP depicting neighborhood vectors and colored by EC neighborhood where each data point is an EC described by its neighborhood composition. **f.** Heatmap showing radial analysis of EC neighborhoods. Neighborhoods were defined by windows containing 5, 10, 30, 50, 100, or 300 cells, and cell type abundance was calculated for each window size. **g.** Stacked bar plot showing cell type composition of the *Cycling TA* neighborhood across window sizes of 5, 10, 30, 50, 100, and 300 cells. **h.** EC neighborhood enrichment per donor normalized to the percentage of ECs in the corresponding tissue subregion. **i.** Area plot of EC neighborhood composition across sections of the small bowel (SB) and colon (CL) averaged across 8 donors. **j.** The percentage of *Immune Enriched* EC neighborhoods is significantly lower throughout the intestine in those who have a history of hypertension (unpaired t-test between donors with and without history of hypertension; p=0.0188). **k.** EC neighborhood composition varies significantly between SB and CL (paired t-test, comparing SB and CL regions for each donor; *: p < 0.05, **: p < 0.01, ***; p < 0.005).

Of the 11 distinct *endothelial cell*-centric neighborhoods, we observed a number of common structures of the intestine such as the follicle and smooth muscle areas (**Fig. 5c**). Indeed, we can see that the neighborhood labels from unsupervised clustering are consistent with the vasculature in these regions, though still are able to discriminate larger from smaller vessels in the same region (**Fig. 5d**). These had similar compositions to several of the neighborhoods that we observed within our original neighborhood analysis^17^, such as *Plasma Cell Enriched* neighborhood. We would expect this as we observe vasculature providing nutrient delivering and absorbing roles throughout the tissue and tissue units.

While we defined these neighborhoods at the scale of 50-cell neighbors, there could be spatial variation in the compositions of these neighborhoods across other numbers of nearest neighbors. To investigate how the cellular composition of neighborhoods varies with spatial scale, we performed a radial analysis using concentric neighborhoods of increasing size, varying from 5, 10, 30, 50, 100 or 300 nearest neighboring cells (**Fig. 5f**). One observation from this analysis is that while *ECs* were present in all neighborhoods, their relative and consistent abundance across the number of nearest neighbors varied.

Based on the variance of the *Cycling TA* neighborhood, we looked specifically into how the percentage of cells surrounding *ECs* changed across the neighbors. Indeed, the relative percentage of *ECs* decreases as the number of nearest neighbors increases (**Fig. 5g**). This pattern may indicate that the *Cycling TA* neighborhood with its enrichment of *transit amplifying cells* and varying *EC* density exhibits a distinct metabolomic demand.

To further investigate the decrease in *EC* enrichment with increasing neighborhood size (**Fig. 5f**), we focused on neighborhoods of size 300 cells. We normalized neighborhood *EC* enrichment to the percentage of *ECs* in the corresponding tissue subregion (mucosa, submucosa, muscularis externa). Tissue-normalized *EC* enrichment revealed that most neighborhoods exhibited higher *EC* enrichment than their subregion average (**Fig. 5h**). These results may indicate that vascular structures are not uniformly distributed throughout each subregion, consistent with varying anatomical demands and the hierarchical branching architecture of the vasculature. The *Smooth Muscle + Innate Immune* neighborhood demonstrated greater *EC* enrichment relative to the muscularis externa average (fold change >1), consistent with increased vascular density (and demand) in these regions.

We previously observed changes in cell neighborhoods across the intestine moving from the duodenum to the descending sigmoid^17^, consequently we hypothesized we would see differences in *EC*-centric neighborhoods as well. We observe changes in several *EC*-centric neighborhoods across the intestine (**Fig. 5i**). Specifically, we see decreases in the *Paneth + Immune* neighborhood which we would expect because *Paneth cells* are largely restricted to the small intestine (**Fig. 5j**). Similarly we see a dramatic decrease in the *Immune Enriched*^57^ neighborhood, while increases in *Cycling TA*, *Stroma*, and *Smooth Muscle + Innate Immune* (**Fig. 5j**). The increase in *Stroma* and *Smooth Muscle + Innate Immune* neighborhoods in the colon may reflect the colon’s denser muscular architecture and its greater demand for oxygen and nutrients. Furthermore, the increase in the *Cycling TA* neighborhood may reflect differences in crypt architecture between the small intestine and colon (large intestine), which likely influence the spatial organization of surrounding vasculature.

We also had previously observed differences in cell type and neighborhood composition associated with metadata of these healthy intestine donors^17^. We expected to see differences in particular with the status of having a history of hypertension or not. Interestingly, significance (p=0.0188) was only found for the *Immune Enriched* neighborhood (**Fig. 5k**). However, there was no correlation of this neighborhood or any *EC* neighborhood percentage with body mass index (BMI). Consequently, this represents a specific shift in the cellular neighborhoods with donors with a history of hypertension. The *Immune Enriched* neighborhood represents vasculature with increased presence of *M1 macrophage* and *CD8+ T cells*, and its decreased frequency in hypertensive donors may reflect either a loss of *CD8+ T cells* preventing the formation of this neighborhood, or reduced *endothelial cell* abundance due to compromised vascular integrity. In conclusion, these results indicate the importance of cell-centric neighborhood analysis and suggest that this approach could be used to find specific neighborhoods/microenvironments from which more specific cell-cell interaction or distance analysis could be completed that may be lost in global averages for an entire tissue.

## Discussion

Various human atlassing efforts^4,6,7,10^ aim to further our understanding of cellular organization and relationships, build comprehensive databases of the adult human blood vasculature^14,15^, and aim to advance our understanding of the angiodiversity and organotypic molecular signatures of human vascular cells across organs^5^.

Spatially resolved omics data enables us to analyze cellular networks—specifically focusing on absolute distances between various cell populations and their nearest endothelial cells—and evaluate the spatial relationships between such cell populations relative to blood vasculature. This can further our understanding of the role of human blood vasculature in spatial organization of cells across organs, tissue regions, and conditions.

This paper focused on open-access methods, data, and tools to measure, visualize, and compare cell distance distributions across organs for spatially resolved omics data. It presented an analysis of cell distance distributions for single-cell spatially resolved omics data for 47M cells across 399 datasets across 12 different tissue types. We processed and compiled a collection of datasets, released publicly, for distance computation and analysis across different regions within a tissue type, different disease conditions in a tissue type, and different samples in tissue types. To enable analysis and comparison, we harmonized the cell type labels across all datasets and defined the labels at three different levels of cell type specificity. Additionally, we spatially registered the datasets, when possible, into the Human Reference Atlas^6^ using the RUI^16^ and the EUI^16^ for posterity and building a harmonized knowledge base of tissue specimen datasets. The cell populations from the spatially registered datasets–currently a subset of 104 datasets that includes skin, spleen, lung, small intestine, and large intestine–are also added to the Human Reference Atlas Cell Type Populations (HRApop) project^58^. Furthermore, we developed and released an open-source web-based tool, Cell Distance Explorer, for interactive analysis and visualization of cell graphs, cell linkages, and distance distributions to enable researchers to see how different conditions might impact cell organization and vascular relationships in tissue samples.

Furthermore, the human oral cavity exhibits a composite tissue identity and spatial organization that varies by anatomical niche. Oral regions blend features of both barrier mucosa (akin to skin, trachea, and esophagus) and glandular tissues (akin to lung and pancreas). Keratinized surfaces such as the gingiva and dorsal tongue resemble skin and esophagus, showing increased epithelial–endothelial distances due to their stratified architecture and adaptation to mechanical stress. In contrast, the parotid and submandibular glands more closely resemble lung and pancreas, where ductal epithelia and secretory acini are densely integrated with vasculature to likely support metabolic output and fluid transport from these specialized cell types. Unlike the intestinal crypt-villus axis, which in this paper exhibits complex *EC*–centered neighborhoods, the oral mucosa lacks such an obvious layering and microarchitectural gradients. Instead, it shows region-specific compartmentalization into distinct barrier and secretory niches that reflect the mouth’s diverse roles. As such, oral tissues occupy a unique position in the body’s spatial landscape, reflected in their distinct patterns of *EC* proximity and subsequently, neighborhood composition.

To capture the nuances between *EC* “neighborhoods,” we investigated the vascular organization across multiple spatial scales by examining how the composition of neighboring cells varies around *ECs* as well as the co-localization of these cellular neighborhoods. Identifying and characterizing these cellular neighborhoods enhances our ability to define the vasculature as an integrated network of functional units within tissue subregions. The results indicate the importance of cell-centric neighborhood analysis and show how this approach can be used to find specific neighborhoods/microenvironments from which more specific cell-cell interaction or distance analysis can be computed.

We find that analyzing spatial single-cell datasets poses a major challenge in terms of cell type label harmonization as different studies may use different cell type naming conventions. Additionally, cell type assignment can be done at varying levels of specificity that increases complexity. While spatially resolved omics datasets have made major advancements, there are still limitations in terms of size and cost of marker panels which limits the number and types of cells that can be detected. These challenges are exacerbated when datasets are compiled from different already published sources and might be ameliorated if a study is defined to use a predefined panel that maximizes cell type overlap across datasets and tissue types. Another challenge for cell distance computation is the fact that analysis in 2D analysis is limited but can be made more accurate if 3D single-cell spatial data is used^53^. While endothelial cell-neighbor distance analysis reveals significant organ-specific biology, determining statistical significance and reliable global differences requires larger sample sizes and formal statistical testing, which we plan to address in future work.

Furthermore, in future work we also plan to extend the analysis beyond euclidean distances and look at cell colocalization and other spatial analyses that can provide additional insights into spatial organization of *endothelial cells*. We also plan to expand the *endothelial cell*-focused neighborhood analysis to other tissue types. Additionally, we hope to collect marker expression information for all datasets and analyze datasets at the biomarker level and not just the cell type level. Last but not least, we are collecting feature requests for the CDE tool to expand current functionality and address evolving user needs.

Taken together, this paper provides an open-access resource (datasets, tools, and analyses) to characterize spatial relationships (specifically, between *endothelial cells* and other cell types using spatially resolved omics data) that helps advance our collective understanding of cell organization across tissues which will, consequently, support the development of various cell atlassing efforts.

## Methods

### Cell Distance Explorer

The Cell Distance Explorer (CDE, https://apps.humanatlas.io/cde) web application was designed, developed, and released in multiple iterations based on user feedback and feature requests. The design process began by synthesizing and prioritizing initial feature requests. Next, we used Figma to start designing the user experience (UX) and user interface (UI). We mocked up an initial design iteration using the Human Reference Atlas Design System, Angular Material development guidelines, and Material 3 design guidelines. Next, we connected the design wireframes to create an interactive, mid-fidelity Figma prototype, to understand how people interacted with the potential app design. We conducted UX research by inviting experts to share candid feedback on their experience using the prototype in moderated, remote usability study sessions. Participants were asked to complete a series of short tasks while interacting with the prototype. This helped us identify areas to improve the design. Additionally, early iterations of the prototype were presented at multiple scientific conferences to capture user needs and user feedback. Both the usability session insights and feedback from the conferences helped us refine the interface design prior to development and implementation. A python package and a lightweight component were also developed and made available for use in Jupyter notebooks and webpages, **see Code Availability**.

### Dataset Collection

**intestine-codex-stanford**^17,20^: This dataset comprises 64 tissue sections (32 from large intestine and 32 from small intestine) from 8 donors. The tissue samples were collected from eight different regions in the intestines (in order of trajectory from the stomach): duodenum, proximal jejunum, mid-jejunum and ileum (four regions in small intestine); ascending, transverse, descending and sigmoid (four regions in large intestine). The dataset uses CODEX imaging technology. All 64 datasets are normal tissue samples. The cell coordinate and cell type annotation data was downloaded from the original published source^17,20^. This study contains two types of endothelial cells at L2 and L3 classification: *endothelial cell* and *endothelial cell of lymphatic vessel*. Since we primarily focus on vascular cells, we use the *endothelial cell* category as the anchor cell type at L2 and L3 classification, while using the *endothelial cell* at L1 classification.

**tonsil-codex-stanford and esophagus-codex-stanford**^51,52^: This dataset comprises two tissue sections, one from tonsil and one from esophagus. The samples were acquired via CODEX imaging. The tonsil dataset is a normal tissue sample and the esophagus dataset is taken from a donor with Barrett’s Esophagus. The cell coordinate and cell type annotation data was downloaded from the original published source. Both datasets contain the *endothelial cell* category at all 3 cell type levels.

**colon-xenium-stanford**^41^: This dataset comprises 29 2D serial tissue sections taken 5 micrometers apart. The sample is taken from a colon polyp. The samples were acquired via Xenium imaging (spatial transcriptomics). The dataset is treated as 29 separate 2D sections in this analysis. The cell coordinate and cell type annotation data was provided by the Stanford team in CSV format and is now published.

**spleen-codex-ufl**^49,50^: This dataset comprises six tissue samples. The samples were acquired via CODEX imaging. The datasets are acquired from normal tissue samples. The cell coordinate and cell type annotation data was directly provided by the JHU team and is published. This dataset contains six 2D datasets and three endothelial cell types at L2 and L3 classification: *endothelial cell of sinusoid*, *endothelial cell of lymphatic vessel*, and *endothelial cell of vascular tree*. We used the *endothelial cell of sinusoid* as the anchor cell due to the much higher cell count across all six samples.

**colon-cycif-sorgerlab**^42^: This dataset comprises 25 five micrometer-thick FFPE serial sections spaced 25 micrometers apart (with the exception of 4 sections that were serially cut) from a donor diagnosed with stage IIIb adenocarcinoma Colorectal Cancer (CRC). The samples underwent ten cycles of CyCIF resulting in 40 channels per plane/section. The cell coordinate and cell type annotation data for each plane was provided by the HMS team.

**lung-codex-urmc**^6^: This dataset comprises two tissue samples from two different donors. The samples were acquired via CODEX imaging. One dataset is a normal sample (D265) and one dataset is a diseased (Bronchopulmonary Disease or BPD) sample (D115). The cell coordinate and cell type annotation data was directly provided by the URMC team. A previous version of the cell coordinates and cell type annotations was published before^6^. The same data was segmented and annotated again by the URMC team for this publication and was made available, **see Data Availability**. This study used two datasets: one normal tissue and one bronchopulmonary disease (BPD) tissue. The dataset contains three types of *ECs* at L2 and L3 classification: *endothelial cell*, *endothelial cell of lymphatic vessel* (only in normal tissue), and *endothelial cell of capillary* (only in BPD tissue), all categorized into *endothelial cell* at L1 classification. At the L2 and L3 classification, *endothelial cell* category is used as the anchor cell. The BPD sample has a comparatively very high *T cell* count, especially *CD4+ alpha-beta T cell* (L3 classification). Some cells in the BPD sample cannot be further disambiguated and hence are labeled as *mixed immune/epithelial cell* population.

**lymphnode-codex-yale**^43–48^: This dataset comprises five tissue samples. The samples were acquired via CODEX imaging. The cell coordinate and cell type annotation data was directly provided by the Yale team and is published. These datasets contain five separate 2D datasets with *endothelial cells* as the anchor cell across all cell type levels. At L1 classification, the *immune cell* population across the five samples is much higher compared to *endothelial cell* and *mesenchymal cell* populations, and across the five samples the *immune cell* population itself varies widely between roughly 1 million to 8 million cells.

**skin-celldive-ge**^53^: This dataset comprises ten 3D tissue samples. The samples were acquired via Cell DIVE imaging. Each of the ten datasets consisted of 24 2D serial sections that were reconstructed into a 3D volume. The cell coordinate and cell type annotation data was downloaded from the original publication. The *endothelial cell* is used as the anchor cell across all three cell type levels.

**skin-confocal-sorgerlab**^39^: This dataset comprises two regions sampled from the same formalin-fixed paraffin embedded (FFPE) specimen and donor (see corresponding manuscript^39^ for clinical data (labelled ‘Dataset 1’). The regions were profiled by CyCIF and imaged by high resolution 3D imaging over 170 z-planes on a laser scanning confocal microscope (Zeiss LSM980 fitted with 40x/1.3NA oil immersion objective lens). The specimen was uniquely cut at ∼25 microns (unlike the traditional 5 microns), thereby allowing analysis of multiple layers of complete cell volumes in the z (vertical) axis. The images were sampled at 140 nm (x,y lateral resolution) and 280 nm (z axial resolution). One dataset is from an invasive margin (i.e., an area of tissue at the edge of a skin cancer where the *tumor cells* have started to invade and spread into surrounding healthy tissue) and the other dataset is from a melanoma in situ region (a much earlier manifestation of melanoma whereby usually diagnosed by the increase in number of *melanocytes* under the epidermis). Both datasets–taken from regions separated by a distance of approximately 1.5 cm along the skin–are true 3D samples prepared and acquired at the same time to reduce batch and operator effects. Datasets were segmented and gated by cell type based on mean intensity of protein expression. The resulting cell coordinates (x, y, z) and cell type annotation data was directly provided by the HMS team and the data is published. They contain *endothelial cell* at all three levels of cell type assignment.

**maternalfetalinterface-mibitof-stanford**^55^: This dataset comprises 209 2D tissue sections from the maternal-fetal interface. The samples were acquired via MIBI-TOF imaging. The data is from normal tissue of 66 individuals. The cell coordinate and cell type annotation data was downloaded from the original published source. We use the *endothelial cell* category as the anchor cell across all three cell type levels. At L1 classification, it has a relatively high *mesenchymal cell* population (mostly *fibroblasts*) across all 209 datasets relative to other L1 classification cell types.

**pancreas-geomx-ufl**^23–34^: This dataset comprises 12 pancreas tissue sections from three donors. The donors represent an 18-year-old male, a 39-year-old female, and a 21 year old female. The tissue samples were collected from four different regions in the pancreas (in order of trajectory): head, neck, body, and tail. The samples were stained using multiplex immunofluorescence and acquired via GeoMx imaging using a 3-plex antibody panel according to published protocol^59^. All 12 datasets are normal tissue samples. The tissue thickness of the samples is 5 µm and the pixel size of the images is 0.4 x 0.4 µm/pixel. The cell coordinate and cell type annotation data was provided by Martha Campbell-Thompson from the UF team. The tissue acquisition and cell segmentation/annotation details are as follows: Whole slide images of pancreas sections were acquired on the NanoString GeoMx Digital Spatial Profiler instrument. These images were exported as OME TIFF files and read into QuPATH version 0.4.1^60^ for analysis. The QuPATH thresholding feature was used to perform a total tissue detection for each pancreas section using an average of all fluorescence channels. This produced a tissue annotation representing combined endocrine and exocrine regions of the pancreas. Next, cell segmentation for each tissue section was performed using the QuPATH positive cell detection feature. Cell detection parameters were optimized for each pancreas section individually due to minor differences in staining intensity between sections. Cell detection of nucleus and cytoplasm was accomplished using the SYTO 83 nuclear stain with 8 µm cell expansion. Cells were classified as positive or negative for each protein marker (insulin, PanCK, CD31/34) using thresholding based on the mean intensity for a given marker across the whole cell. Cell detections for each pancreas section were exported from QuPATH as CSV files containing +/- values for each protein marker as well as 2D cell coordinates in µm. R scripts were used to format the CSV files appropriately for downstream analysis. In the pancreas-geomx-ufl datasets, the *unknown cells* are most abundant but contain mostly *acinar cells* following aspiration of *beta cells*, *duct cells*, and *ECs* by the DSP instrument. Since this data was not published before as part of a previous study, it is published and made available as part of this paper, **see Data Availability**. This study contains four cell type categories at all three cell type levels: *endothelial cells*, *ductal epithelial cells* (*epithelial cell* at L1 classification and *gland epithelium cell* at L2 classification), *pancreatic beta cells* (*beta cell* at L2 classification and *epithelial cell* at L1 classification), and *unknown cells*. The *pancreatic beta cell* population is very low compared to other cell types, as expected due to scarcity of islets in general, and the majority of cells are labeled as *unknown cells* of which *acinar cells* are the large majority.

**oralcavity-codex-czi**^35–37^: This dataset comprises 13 tissue sections from the mouth (also called oral cavity). The tissue samples were collected from 5 different regions in the mouth: Buccal Mucosa and Minor Salivary Glands (two datasets), Gingiva (two datasets), Parotid (four datasets), Submandibular (two datasets), and Tongue (three datasets). The samples were acquired via PhenoCycler Fusion imaging. All 13 datasets are normal tissue samples. The cell coordinate and cell type annotation data was provided by the Virginian Commonwealth University team and is published. This study contains two types of *ECs* at L3 classification: *endothelial cell of vascular tree* and *endothelial cell of lymphatic vessel*, both of which are categorized similarly at L2 classification, and as *endothelial cells* at L1 classification. At L2 and L3 classification, the *endothelial cell of vascular tree* is used as the anchor cell.

**bonemarrow-codex-chop**^40^: This dataset comprises 20 tissue sections from the bone marrow in the head of femur: 12 from normal (NBM) samples, 5 from acute myeloid leukemia (AML), and 3 from Negative lymphoma staging bone marrow biopsies (NSM). The samples were acquired via CODEX imaging. The cell coordinate and cell type annotation data was downloaded from the original published source. At L2 and L3 classification, they contain two categories of *ECs*: *endothelial cell of artery* and *endothelial cell of sinusoid*. We use the latter as the anchor cell due to the relatively higher cell counts across all samples.

### Data Preprocessing

All collected datasets were processed to standardize the file formats and schema. The cell coordinates (x, y, and z if 3D) were extracted from the original data sources along with the assigned cell type labels and were saved as a CSV file. A single CSV file per individual dataset was created. When relevant, the cell coordinates were multiplied by the pixel size to ensure all coordinates were in micrometers. In the lung dataset, the relevant data was extracted from a H5ad file format and saved as a CSV. Bone marrow data was extracted from an R object file and saved as CSV. For pancreas data, cell type labels were assigned based on marker positivity/negativity. Where relevant and available, datasets were also divided into unique regions within organs, separated by condition (disease), and/or donors. The original and processed data is made available, see **Data Availability**.

### Cell Type Harmonization

The original cell type labels across all datasets were mapped to Level Three (L3, most specific), Level Two (L2), and Level One (L1, most general) cell types based on expert-validated mappings. The original cell type information is retained for posterity. All cell type labels across the three levels were also mapped to Cell Ontology; the Cell Ontology label (CL label) and Cell Ontology ID (CL ID) are added to the dataset files. When mapping the cell type labels to Cell Ontology labels, we also define a match level (exactMatch or narrowMatch) depending on how close the label in the dataset matches the Cell Ontology typology. This process was done manually and iteratively, where ontology experts defined the appropriate CL label and CL ID for each cell type and domain experts (computational biologists and clinicians who provided the datasets) defined and reviewed the cell type labels across all three levels. The labels were defined to maximize the overlap between datasets, and experts and data providers made the final call in all cases, especially when insufficient information/markers were available for the dataset.

The entire final data collection, including all cell type labels (Original, L1, L2, L3) and CL labels and CL IDs are publicly available, **see Data Availability**. The code used to process the datasets is also available, **see Code Availability**. The complete listing of all cell types, including original cell type labels and harmonized cell type labels, is available at https://github.com/cns-iu/hra-cell-distance-analysis/blob/main/data/mapping_files/generated_cell_type_complete_crosswalk.csv.

### Registration User Interface and Exploration User Interface

The size, location, and rotation of all tissue specimen data was registered using the HRA RUI^16^. The registration process required close coordination between data providers and the registration coordinator following standard operating procedures^61^ to create and position 3D virtual tissue blocks that match the physical tissue block specimens in terms of size, location, and orientation. The registered data was then federated in the HRA knowledge base and all registered data was made available in the general Exploration User Interface (EUI)^16^ and in an EUI instance specific to the collection used in this meta-study (https://hubmapconsortium.github.io/hra-registrations/vccf-federated/). The virtual blocks were enriched with additional metadata, including information about the donor, assay types, and associated publications. Data providers reviewed and verified the accuracy of the registered information before final publication in the Exploration User Interface. For 243 datasets from the maternal-fetal interface, bone marrow, tonsil, and oral cavity, no 3D reference organ was available; for 1 dataset from the esophagus, no location information existed; hence those 244 datasets were not registered.

### Distance Computations

For all 399 datasets in the collection, we compute cell-to-nearest-anchor-cell distances within a defined maximum distance threshold. In this paper, *vascular endothelial cell* types (or subtypes) are used as the anchor cells. The distance is based on euclidean distance and a maximum distance threshold of 200 µm is used for all datasets. The input file, called nodes.csv, has 4 columns with values for x, y, and z position of cells and the name of cells, one row per cell; the z position is optional. The resulting output file, edges.csv, lists the computed linkages and euclidean distance values from each non-anchor node to the closest anchor node. We use an optimized python script for computing the distances which is much faster than the naive approach. The script is made available publicly, **see Code Availability**. A JavaScript version of the script is used in the Cell Distance Explorer to do similar computations in the web browser. All processed edges.csv files are made available, **see Data Availability**.

Coefficient of Variability (CV) is a statistical measure of dispersion of a distribution i.e., it shows the extent of variability in relation to the mean of the population which shows what cell types or regions have a higher relative variability regardless of their absolute distances. It is defined as the ratio of the standard deviation to the mean of the data. A CV of 0 indicates no variation, while higher CV values signify greater variability in the data.

### Endothelial cell-focused neighborhood analysis

We initially generated 20 clusters, these clusters were then assessed for both cellular composition and spatial localization. Clusters that were spatially co-localized and exhibited similar cell type compositions were subsequently merged, resulting in 11 distinct endothelial cell–centric neighborhoods.

*Endothelial cell (EC)* enrichment was calculated for each cell neighborhood as the proportion of *ECs* among total cells in the neighborhood. To account for differences in baseline *EC* abundance across tissue subregions and between donors, *EC* enrichment was normalized per donor by dividing the neighborhood *EC* enrichment by the mean *EC* enrichment of the corresponding tissue subregion (mucosa, submucosa, or muscularis externa), see **Fig. 5b**.

### Neighborhood analysis with donor metadata

*EC* neighborhood percentage was calculated for each image/region by finding the proportion of cells that belonged to each neighborhood. For all statistical tests, replicates were defined by individual donors. To investigate correlations with donor BMI, the neighborhood percentages for each donor were averaged, and a linear regression was performed between BMI and the average neighborhood percentage per donor.

For assessing the effect of history of hypertension, *EC* neighborhood percentage was similarly calculated per donor by averaging across all regions per donor. For each neighborhood, an unpaired t-test was conducted between the hypertension and no hypertension groups.

For investigating the difference in *EC* neighborhood composition between the small bowel (SB, also called small intestine) and colon (CL, also called large intestine), the neighborhood percentages were first averaged across regions of the SB or CL for each donor, resulting in data pairs of neighborhood percentages in the SB and CL per donor. For each neighborhood, a paired t-test was conducted between the SB and CL neighborhood percentages.

## Acknowledgements

We thank Tracey Lynn Theriault for the final figure design. We thank the authors of the original studies and datasets that have been used in this meta-analysis for their contributions to the respective publications. This work is funded by the NIH Common Fund through the Office of Strategic Coordination/Office of the NIH Director under awards OT2OD033756 (K.B.), OT2OD026671 (K.B.), and OT2OD030545 (K.B.); by the Cellular Senescence Network (SenNet) Consortium through the Consortium Organization and Data Coordinating Center (CODCC) under award number U24CA268108 (K.B.); by the Kidney Precision Medicine Project grant U2CDK114886 (K.B.); by the NIDDK under awards U24DK135157 (K.B.) and U01DK133090 (K.B.); by the NIH Common Fund through the Office of Strategic Coordination/Office of the NIH Director under awards 3OT2OD033759-01S4 and 3U54AG07593 (JWH); by the ADA Science & Research Institute (startup funds & Volpe Research Scholar Award, K.M.B.); and the Department of Oral and Molecular Craniofacial Biology, Philips Institute for Oral Health Research (startup funds, K.M.B.); VCU Massey Cancer Center, National Council for Scientific and Technological Development; CZI Pediatric Networks for the Human Cell Atlas, “Mapping the Pediatric Inhalation Interface.” Grant Number: 2021-237918 (K.M.B.); and by grant awards U54HL165443 (GSP), U01HL148861 (GSP), U54DK127823 (MCT), CCBIR grants U54-CA268072, U2C-CA233280, U2C-CA233262 (CY), OT2OD033760 (JF), HTAN grant award U2CCA233311 (CZ, EMM), OT2OD026675 (A.B. as NIH JumpStart Award), OT2OD033759 (A.B. as NIH JumpStart Fellowship), 1R03OD039970-01 (AB, BWH), U54CA274509 (RF), UH3CA257393 (RF), RF1MH128876 (RF), U54AG079759 (RF), U54AG076043 (RF), U01CA294514 (RF), R01CA245313 (RF), RM1MH132648 (RF), U54AR081775 (FG, AK).

## Data Availability

All datasets (cell coordinates and cell types) used in the paper, original and processed (including harmonized cell type labels), are made publicly available (https://cns-iu.github.io/hra-cell-distance-analysis/). For more information and original imaging datasets for individual studies used in this paper, see relevant citations in the **Methods** section. **Pancreas dataset** (GeoMX) is available on Figshare (https://figshare.com/projects/HuBMAP_TMC_-_Pacific_Northwest_National_Laboratory_GeoMX_DSP_Images/256367).

## Code Availability

All code for data processing and analysis for **distance analysis** is available on GitHub (https://github.com/cns-iu/hra-cell-distance-analysis). Rendered Jupyter notebooks for the entire **workflow** are also available on GitHub Pages (https://cns-iu.github.io/hra-cell-distance-analysis). The file containing all cell types, original and **3-level typology**, mapped to **Cell Ontology,** is available as a CSV file on GitHub (https://github.com/cns-iu/hra-cell-distance-analysis/blob/main/data/mapping_files/generated_cell_type_complete_crosswalk.csv).

Code for **hierarchical neighborhood analysis** is available on GitHub (https://github.com/HickeyLab/Vasculature_neighborhoods). The **Cell Distance Explorer** application is available on the Human Reference Atlas website (https://apps.humanatlas.io/cde). More information, including **usage tutorial**, about the CDE can be found at https://humanatlas.io/user-story/5. The **python package** for CDE can be found as part of HRA Jupyter Widgets (https://github.com/x-atlas-consortia/hra-jupyter-widgets/). Documentation to **embed the CDE in webpages** as a lightweight component is available at https://github.com/hubmapconsortium/hra-ui/blob/main/apps/cde-visualization-wc/EMBEDDING.md.

## Author contributions

**YJ** led the data collection, data processing, data analysis, and data visualization for distance analysis, primary contributor to the design and implementation of the study, aided the interpretation of the data and results, co-wrote the paper; **JJ** and **RC** contributed to the neighborhood analysis, data visualization, results/data interpretation, and contributed to writing and revising of manuscript; **APB** and **EMQ** mapped all cell types to Cell Ontology; **EMQ** also tested Cell Distance Explorer web application and contributed to writing the manuscript; **EM** served as the user experience product designer for the Cell Distance Explorer web application, conducted hands-on usability testing and refined the interface through continuous collaboration with both end users and project contributors; **BWH** led the development of the Cell Distance Explorer application; **DQ** coordinated the spatial registration of tissue samples into the Human Reference Atlas; **SLE** performed single cell analyses using QuPATH on GeoMX MxIF-stained FFPE sections for pancreas data; **AE** and **NF** performed CODEX multiplexed immunofluorescence staining on FFPE lymph node tissue sections; **AB** utilizes datasets from this effort and integrates them into the HRA Cell Type Population (HRApop) effort; **QTE** created OMAPs, ran original experiments, supported cell annotations, analysis, and 3D integration for the oral cavity data; **BM** generated and cell assigned all the oral cavity tissues, and contributes to the HRA; **CZ** did the Xenium data analysis for intestine; **EMM** did primary project management for xenium intestine data; **JMP** did MxIF imaging and analysis for lung data; **MJ** did image segmentation and clustering of MxIF lung data; **RSM** did experimental design and MxIF image analysis for lung data; **RF** led the study design, supervision, and data interpretation for the lymph node dataset; **JF** contributed annotated CODEX data for spleen; **FG** contributed skin CELLDIVE data; **AK** contributed skin CELLDIVE data; **CY** distributed 3D CyCIF colorectal cancer and skin melanoma datasets, for skin melanoma dataset **CY** performed experiment design, 3D data and image acquisition, 3D image processing, single cell image quantification, cell gating, interpretation of results, and contributed to writing of manuscript; **MCT** did MxIF staining for GeoMX DSP whole transcriptome assay oversight, single cell analysis oversight of MxIF stained sections, interpretation of results, writing/editing manuscript, lead contact pancreas ASCT+B table, FTU design, and 3D pancreas for Human Reference Atlas; **GP** led the study design, supervision, and data interpretation for the lung dataset, conceptualized the 3-level cell type hierarchy for the paper, major contributor to the cell type harmonization effort; **KMB** created OMAPs, ran original experiments, supported cell annotations, analysis, and 3D integration for oral cavity datasets; **JWH** conceptualized multiscale analysis, integration with cell-cell distance analysis, oversee analysis and interpretation, neighborhood analysis, data visualization, results/data interpretation, and contributed to writing and revising of manuscript; **KB** leads the Human Reference Atlas and contributes to vasculature-based Common Coordinate System (VCCF) construction, conceptualized the paper and contributed to writing and revising of manuscript.

## Supplementary Tables

**Supplementary Table 1.**
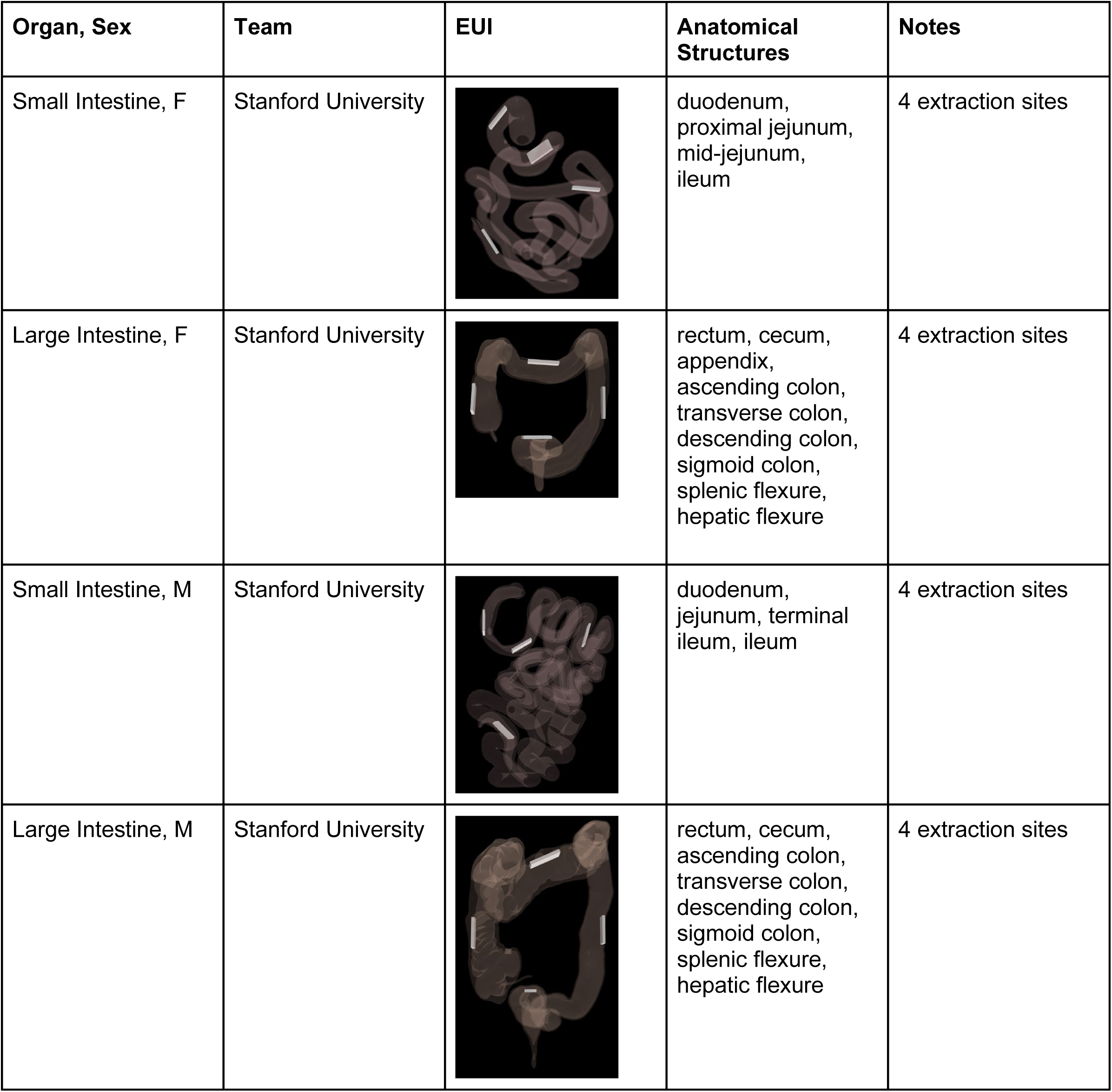
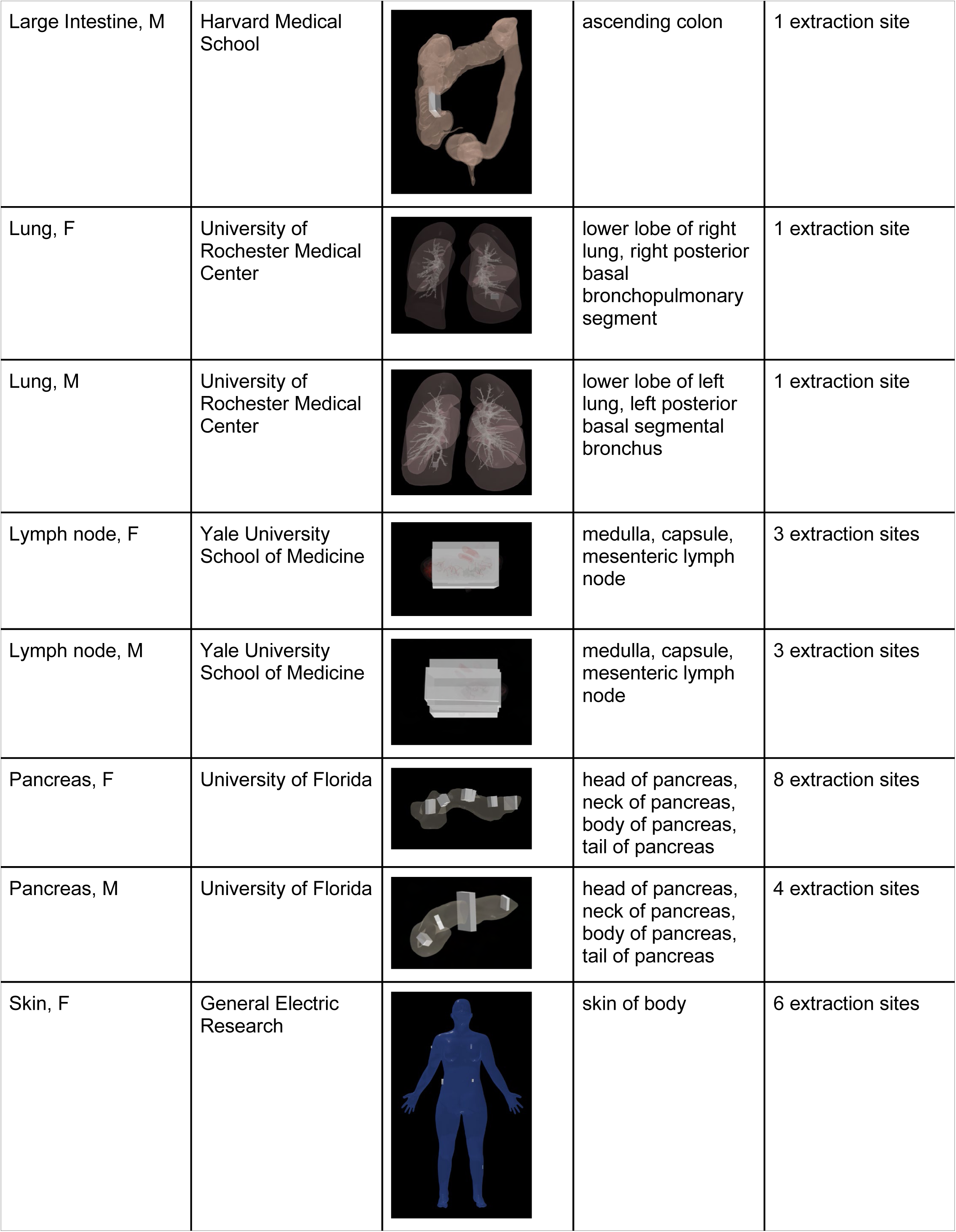
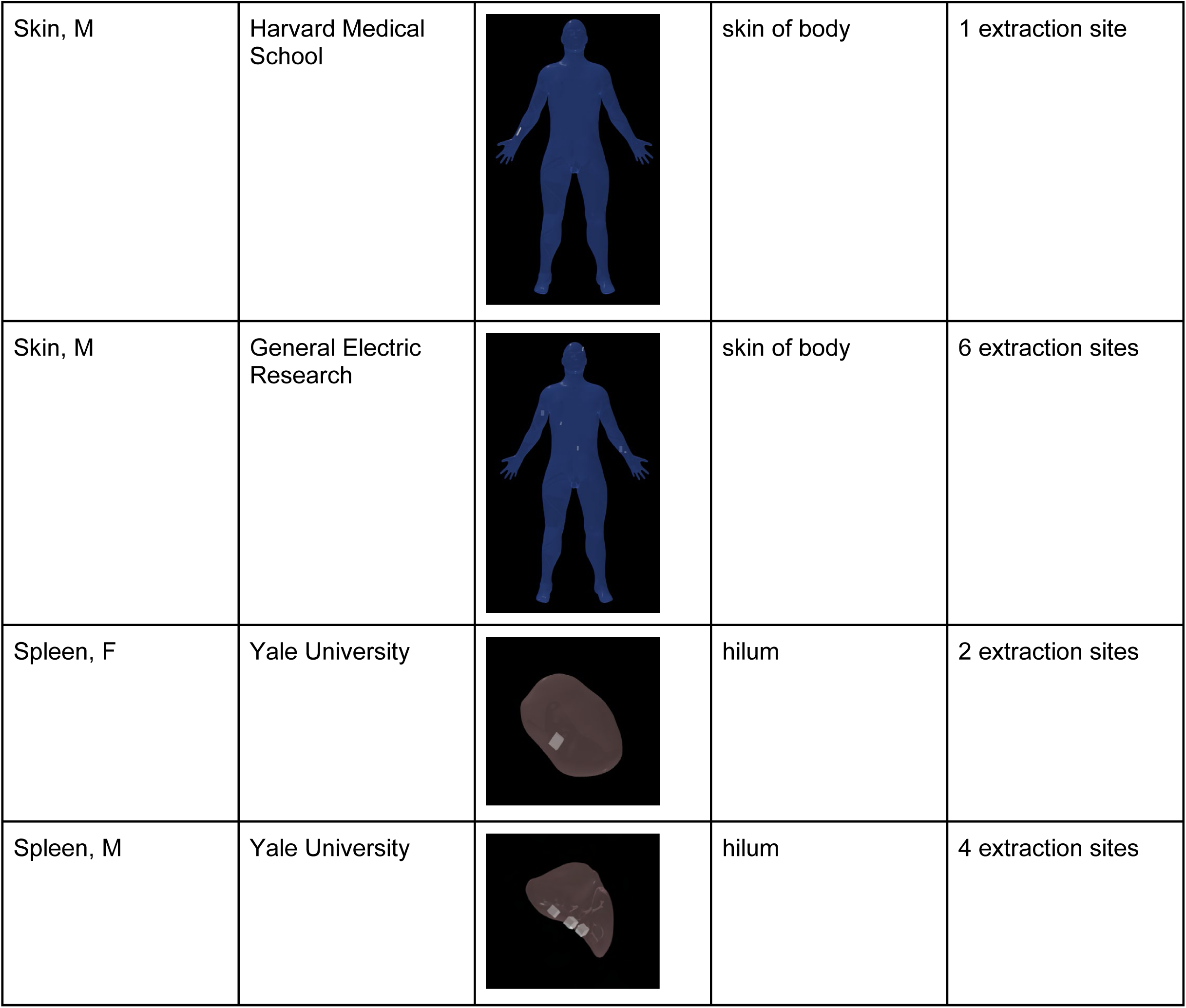
Screenshots of individual EUI organs for studies registered using the RUI.

| Organ, Sex | Team | EUI | Anatomical Structures | Notes |
| --- | --- | --- | --- | --- |
| Small Intestine, F | Stanford University |  | duodenum, proximal jejunum, mid-jejunum, ileum | 4 extraction sites |
| Large Intestine, F | Stanford University |  | rectum, cecum, appendix, ascending colon, transverse colon, descending colon, sigmoid colon, splenic flexure, hepatic flexure | 4 extraction sites |
| Small Intestine, M | Stanford University |  | duodenum, jejunum, terminal ileum, ileum | 4 extraction sites |
| Large Intestine, M | Stanford University |  | rectum, cecum, ascending colon, transverse colon, descending colon, sigmoid colon, splenic flexure, hepatic flexure | 4 extraction sites |
| Large Intestine, M | Harvard Medical School |  | ascending colon | 1 extraction site |
| Lung, F | University of Rochester Medical Center |  | lower lobe of right lung, right posterior basal bronchopulmonary segment | 1 extraction site |
| Lung, M | University of Rochester Medical Center |  | lower lobe of left lung, left posterior basal segmental bronchus | 1 extraction site |
| Lymph node, F | Yale University School of Medicine |  | medulla, capsule, mesenteric lymph node | 3 extraction sites |
| Lymph node, M | Yale University School of Medicine |  | medulla, capsule, mesenteric lymph node | 3 extraction sites |
| Pancreas, F | University of Florida |  | head of pancreas, neck of pancreas, body of pancreas, tail of pancreas | 8 extraction sites |
| Pancreas, M | University of Florida |  | head of pancreas, neck of pancreas, body of pancreas, tail of pancreas | 4 extraction sites |
| Skin, F | General Electric Research |  | skin of body | 6 extraction sites |
| Skin, M | Harvard Medical School |  | skin of body | 1 extraction site |
| Skin, M | General Electric Research |  | skin of body | 6 extraction sites |
| Spleen, F | Yale University |  | hilum | 2 extraction sites |
| Spleen, M | Yale University |  | hilum | 4 extraction sites |

## References

1. Acharya, B. R., Chavkin, N. W. & Hirschi, K. K. Chapter 1 - Development of, and environmental impact on, endothelial cell diversity. in The Vasculome (ed. Galis, Z. S.) 5–15 (Academic Press, 2022). doi:10.1016/B978-0-12-822546-2.00013-7.

2. Bidanta, S. et al. Functional tissue units in the Human Reference Atlas. Nat. Commun. 16, 1526 (2025).

3. Marx, V. Atlases galore: where to next? Nat. Methods 21, 2203–2208 (2024).

4. Jain, S. et al. Advances and prospects for the Human BioMolecular Atlas Program (HuBMAP). Nat. Cell Biol. 25, 1089–1100 (2023).

5. Barnett, S. N. et al. An organotypic atlas of human vascular cells. Nat. Med. 30, 3468–3481 (2024).

6. Börner, K. et al. Human BioMolecular Atlas Program (HuBMAP): 3D Human Reference Atlas construction and usage. Nat. Methods 22, 845–860 (2025).

7. Lee, P. J. et al. NIH SenNet Consortium to map senescent cells throughout the human lifespan to understand physiological health. *Nat*. Aging 2, 1090–1100 (2022).

8. Suryadevara, V. et al. SenNet recommendations for detecting senescent cells in different tissues. Nat. Rev. Mol. Cell Biol. 25, 1001–1023 (2024).

9. de Bruijn, I. et al. Sharing data from the Human Tumor Atlas Network through standards, infrastructure and community engagement. Nat. Methods 22, 664–671 (2025).

10. Rozenblatt-Rosen, O. et al. The Human Tumor Atlas Network: Charting Tumor Transitions across Space and Time at Single-Cell Resolution. Cell 181, 236–249 (2020).

11. Bodenmiller, B. Highly multiplexed imaging in the omics era: understanding tissue structures in health and disease. Nat. Methods 21, 2209–2211 (2024).

12. Method of the Year 2024: spatial proteomics. Nat. Methods 21, 2195–2196 (2024).

13. Bussi, Y. & Keren, L. Multiplexed image analysis: what have we achieved and where are we headed? Nat. Methods 21, 2212–2215 (2024).

14. Weber, G. M., Ju, Y. & Börner, K. Considerations for Using the Vasculature as a Coordinate System to Map All the Cells in the Human Body. Front. Cardiovasc. Med. 7, (2020).

15. Boppana, A. et al. Anatomical structures, cell types, and biomarkers of the healthy human blood vasculature. Sci. Data 10, 452 (2023).

16. Börner, K. et al. Tissue registration and exploration user interfaces in support of a human reference atlas. *Commun*. Biol. 5, 1–9 (2022).

17. Hickey, J. W. et al. Organization of the human intestine at single-cell resolution. Nature 619, 572–584 (2023).

18. Diehl, A. D. et al. The Cell Ontology 2016: enhanced content, modularization, and ontology interoperability. J. Biomed. Semant. 7, 44 (2016).

19. CL - Ontology Lookup Service. https://www.ebi.ac.uk/ols4/ontologies/cl.

20. Hickey, J. Processed single cell data from CODEX multiplexed imaging of the human intestine. 5864694150 bytes Dryad 10.5061/DRYAD.PK0P2NGRF (2022).

21. Matusiak, M. et al. Spatially Segregated Macrophage Populations Predict Distinct Outcomes in Colon Cancer. Cancer Discov. 14, 1418–1439 (2024).

22. Bernier-Latmani, J., González-Loyola, A. & Petrova, T. V. Mechanisms and functions of intestinal vascular specialization. J. Exp. Med. 221, e20222008 (2023).

23. Kates, H. & Campbell-Thompson, M. OME TIFF image GeoMX whole transcriptome assay from P2-4AD. figshare 10.6084/m9.figshare.29661272.v4 (2025).

24. Campbell-Thompson, M. & Kates, H. OME TIFF image GeoMX whole transcriptome assay P2-13. figshare 10.6084/m9.figshare.29669540.v4 (2025).

25. Campbell-Thompson, M. & Kates, H. OME TIFF image GeoMX whole transcriptome assay P2-19. figshare 10.6084/m9.figshare.29669573.v2 (2025).

26. Campbell-Thompson, M. & Kates, H. OME TIFF image GeoMX whole transcriptome assay P3-3A. figshare 10.6084/m9.figshare.29669618.v1 (2025).

27. Campbell-Thompson, M. & Kates, H. OME TIFF image GeoMX whole transcriptome assay P3-7A. figshare 10.6084/m9.figshare.29669648.v1 (2025).

28. Campbell-Thompson, M. & Kates, H. OME TIFF image GeoMX whole transcriptome assay P3-9A. figshare 10.6084/m9.figshare.29669675.v1 (2025).

29. Campbell-Thompson, M. & Kates, H. OME TIFF image GeoMX whole transcriptome assay P3-13A. figshare 10.6084/m9.figshare.29669726.v1 (2025).

30. Campbell-Thompson, M. & Kates, H. OME TIFF image GeoMX whole transcriptome assay P4-3A. figshare 10.6084/m9.figshare.29669789.v3 (2025).

31. Campbell-Thompson, M. & Kates, H. OME TIFF image GeoMX whole transcriptome assay P4-6A. figshare 10.6084/m9.figshare.29669831.v1 (2025).

32. Campbell-Thompson, M. & Kates, H. OME TIFF image GeoMX whole transcriptome assay P4-12A. figshare 10.6084/m9.figshare.29669861.v1 (2025).

33. Campbell-Thompson, M. & Kates, H. OME TIFF image GeoMX whole transcriptome assay P4-22. figshare 10.6084/m9.figshare.29669882.v2 (2025).

34. Campbell-Thompson, M. & Kates, H. OME-TIFF image GeoMX whole transcriptome assay from P2-7A. figshare 10.6084/m9.figshare.29661308.v2 (2025).

35. Easter, Q. T. et al. Single-cell and spatially resolved interactomics of tooth-associated keratinocytes in periodontitis. Nat. Commun. 15, 5016 (2024).

36. Matuck, B. F. et al. The Immunoregulatory Architecture of the Adult Oral Cavity. 2024.12.01.626279 Preprint at 10.1101/2024.12.01.626279 (2024).

37. Huynh, K. L. A. et al. Deconvolution of cell types and states in spatial multiomics utilizing TACIT. Nat. Commun. 16, 3747 (2025).

38. Pranzatelli, T. J. F. et al. GZMK+CD8+ T cells Target A Specific Acinar Cell Type in Sjögren’s Disease. *Res. Sq.* rs.3.rs-3601404 (2024) doi:10.21203/rs.3.rs-3601404/v2.

39. Yapp, C. et al. Highly Multiplexed 3D Profiling of Cell States and Immune Niches in Human Tumours. 2023.11.10.566670 Preprint at 10.1101/2023.11.10.566670 (2025).

40. Bandyopadhyay, S. et al. Mapping the cellular biogeography of human bone marrow niches using single-cell transcriptomics and proteomic imaging. Cell 187, 3120–3140.e29 (2024).

41. NCI Human Tumor Atlas Network. https://data.humantumoratlas.org/publications/hta10_2025_tbd_rongduo-han.

42. Lin, J.-R. et al. Multiplexed 3D atlas of state transitions and immune interaction in colorectal cancer. Cell 186, 363–381.e19 (2023).

43 Enninful, A., Fan, R., Farzad, N. & Zhong, M. PhenoCycler data from the lymph node of a 22.0-year-old unknown female. SenNet Consortium 10.60586/SNT228.XBSK.467 (2025).

44. Enninful, A., Fan, R., Farzad, N. & Zhong, M. PhenoCycler data from the lymph node of a 86.0-year-old white male. SenNet Consortium 10.60586/SNT832.RKFV.643 (2025).

45. Enninful, A., Fan, R., Farzad, N. & Zhong, M. SNT222.DNBF.782 | Dataset | SenNet. 10.60586/SNT222.DNBF.782 (2025).

46. Enninful, A., Fan, R., Farzad, N. & Zhong, M. SNT325.JRSJ.469 | Dataset | SenNet. 10.60586/SNT325.JRSJ.469 (2025).

47. Enninful, A., Fan, R., Farzad, N. & Zhong, M. SNT584.CWSK.568 | Dataset | SenNet. 10.60586/SNT584.CWSK.568 (2025).

48. Enninful, A., Fan, R., Farzad, N. & Zhong, M. SNT625.XDTN.459 | Dataset | SenNet. 10.60586/SNT625.XDTN.459 (2025).

49. dos Santos Peixoto, R., et al. Characterizing cell-type spatial relationships across length scales in spatially resolved omics data. Nat. Commun. 16, 350 (2025).

50. Currlin, S. et al. Immune, endothelial and neuronal network map in human lymph node and spleen. 2021.10.20.465151 Preprint at 10.1101/2021.10.20.465151 (2022).

51. Brbić, M. et al. Annotation of spatially resolved single-cell data with STELLAR. Nat. Methods 19, 1411–1418 (2022).

52. Hickey, J. CODEX multiplexed imaging cell datasets used for using STELLAR to transfer cell type annotations to other tissues and donors. 998655507 bytes Dryad 10.5061/DRYAD.G4F4QRFRC (2022).

53. Ghose, S. et al. 3D reconstruction of skin and spatial mapping of immune cell density, vascular distance and effects of sun exposure and aging. *Commun*. Biol. 6, 718 (2023).

54. Wang, X.-N. et al. A Three-Dimensional Atlas of Human Dermal Leukocytes, Lymphatics, and Blood Vessels. J. Invest. Dermatol. 134, 965–974 (2014).

55. Greenbaum, S. et al. A spatially resolved timeline of the human maternal–fetal interface. Nature 619, 595–605 (2023).

56. Hickey, J. W. et al. T cell-mediated curation and restructuring of tumor tissue coordinates an effective immune response. Cell Rep. 42, 113494 (2023).

57. Brown, H. & Esterházy, D. Intestinal immune compartmentalization: implications of tissue specific determinants in health and disease. Mucosal Immunol. 14, 1259–1270 (2021).

58. Bueckle, A. et al. Constructing and Using Cell Type Populations of the Human Reference Atlas. 2025.08.14.670406 Preprint at 10.1101/2025.08.14.670406 (2025).

59. Fu, A. et al. GeoMX Digital Spatial Profiler Whole Transcriptome Assay for Human Pancreas. (2022).

60. Bankhead, P. et al. QuPath: Open source software for digital pathology image analysis. Sci. Rep. 7, 16878 (2017).

61. Bueckle, A. & Qaurooni Fard, D. Using the Standalone Registration User Interface. (2024).

